# A genomic and functional framework for the rapid domestication of the wild plant *Chenopodium album*

**DOI:** 10.64898/2026.01.13.699046

**Authors:** Sanskriti Vats, Alexandra Sanfeliu Meliá, Kate Escobar, Jan Günther, Mariela González-Ramírez, Beatriz Cavaleiro César, Leen Leus, Poul Erik Jensen, Katrijn Van Laere, Søren Bak, Pablo D. Cárdenas

## Abstract

Global reliance on a small number of genetically uniform crops makes our food system increasingly vulnerable to pests, diseases, and climate change, highlighting the need to develop resilient local species as crops. *Chenopodium album*, a stress-tolerant, protein-rich wild plant whose seeds were part of prehistoric Northern European diets and whose leaves are still foraged worldwide, remains undomesticated despite its agrifood potential. We established a Danish collection of 143 accessions and combined seed metabolomics, ploidy assessment and genomics to uncover the molecular basis of key nutritional and anti-nutritional traits. Seed profiling revealed substantial variation in protein content (14-22%), comparable to or higher than major crops, and 16 distinct triterpenoid saponins, which are widespread bitter and anti-nutritional compounds. Seed production of field-grown lines reached up to 1.5 t/ha in trials conducted in Denmark, demonstrating promising yield potential. A high-quality tetraploid genome of a low-saponin line was assembled and contrasted with resequencing of a diploid high-saponin line in order to uncover the genetic basis of saponin variation in *C. album*. Comparative genomic, phylogenetic, and transcriptomic analyses identified structural variants and candidate genes associated with saponin biosynthesis, and functional validation confirmed the coordinated activity of a β-amyrin synthase, three CYP716 cytochromes P450, and a glucuronosyltransferase that reconstitute the core *C. album* saponin pathway. Together, these results define the genomic and biochemical foundation of *C. album*, establishing a platform for its rapid domestication as a locally adapted, high-protein seed crop and a model for translating wild plant diversity into future food security.

## Introduction

Our global agrifood system relies heavily on a narrow range of uniform crops that supply most of the world’s caloric intake. Rice, wheat, maize, and potato together provide nearly 60% of the global food energy, despite the existence of more than 30,000 edible plant species (Li and Siddique, 2018). Such dependence on a few staples has improved productivity but left food systems vulnerable to pests, diseases, and the growing impacts of climate change (Deutsch et al., 2018). As soils degrade, weather extremes increase, and geopolitical conflicts intensify, the stability of globalized supply chains is threatened, underscoring the need for resilient, locally adapted crops.

Harnessing local plant diversity offers a practical path to enhanced food security. Indigenous species thrive under minimal inputs in their native environments, while non-native crops require intensive inputs and costly adaptation efforts to new conditions (Shelef et al., 2017). Modern genomic and breeding tools now enable the domestication of wild plants, allowing local biodiversity to be translated into agronomic and nutritional value. Diversifying agriculture with such species can reduce production risks and enhance both ecological and dietary resilience.

Among these overlooked wild species, *Chenopodium album* (white goosefoot) is particularly promising. This stress-tolerant annual was partly domesticated in the Himalayas (Partap and Kapoor, 1985) and consumed for its seeds in prehistoric diets in present-day Denmark (Stokes and Rowley-Conwy, 2002). Today it is largely regarded as a weed, although it is still foraged for its edible leaves in countries such as India and Nepal. *C. album* exhibits strong crop potential due to its high nutritional value, strong adaptability to marginal soils, and resistance to pests and diseases (Bajwa et al., 2019; Bhardwaj et al., 2023; Ghazaryan et al., 2024), positioning it as a potential local crop for temperate regions.

In addition to the limited knowledge of the agronomic performance of *C. album*, a key challenge for its domestication is the presence of triterpenoid saponins, specialized metabolites that contribute bitterness and reduce nutrient digestibility (Cárdenas et al., 2019). Although these compounds serve protective roles against herbivores and pathogens, their accumulation in seeds restricts palatability and market potential. Until now, the molecular basis of saponin biosynthesis in *C. album* has remained unresolved, and dedicated omics resources to support genetic and metabolic studies have been lacking.

To address these gaps and illustrate the potential for domestication of local wild species as future food crops, we established a collection of 143 *C. album* accessions from across Denmark and characterized their seed protein content and saponin profiles. Two contrasting lines, one high and one low in saponin accumulation, were selected for agronomic performance and molecular analyses. We generated a high-quality annotated tetraploid genome assembly for the low-saponin line and resequenced the high-saponin line, integrated transcriptomic and metabolomic data, and identified candidate genes involved in saponin biosynthesis. Functional assays confirmed core enzymatic steps in the pathway leading to oleanolic acid-derived saponins.

Together, these omics resources provide the foundation for the rapid domestication of *C. album*. We demonstrate how a wild, resilient, and protein-rich species can be transformed into a promising local food crop through the integration of phenotyping, omics analyses, and functional validation. By bridging a key gap in the understanding of saponin biosynthesis, this work supports the broader goal of harnessing wild plant diversity for sustainable and climate-resilient food systems.

## Results and Discussion

### Natural variation in protein and saponin content among Danish *C. album* lines

We established and phenotyped a collection of 143 *C. album* lines from 16 locations across Denmark, revealing extensive, previously undocumented variation in seed protein and saponin content. All lines were advanced by four generations of self-fertilization to reduce within-line heterogeneity. Seed protein content ranged from 14% to 22% (Figure 1), demonstrating substantial nutritional diversity and potential for selecting high-protein genotypes, consistent with trends in other Amaranthaceae crops (Ruiz et al., 2017).

**Figure 1.**
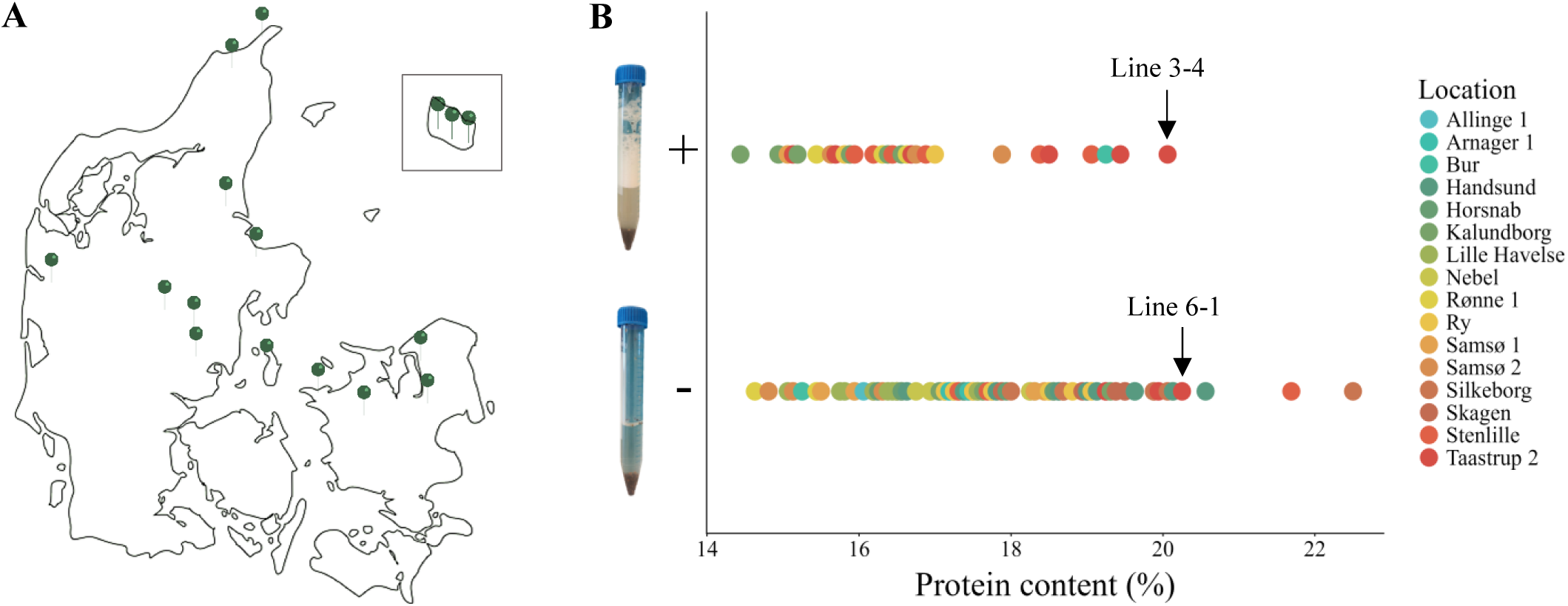
Geographic origin and seed trait diversity of Danish *Chenopodium album* accessions. **A.** Map of Denmark showing the 16 locations where 143 wild *C. album* accessions were collected. **B**. Seed protein and saponin content across the 143 accessions. Saponin content was estimated using a foaming assay. Each point represents one accession and is colored by collection location. Overall, 32 lines showed high levels of saponins (+), while the remaining 111 lines showed low levels (−). Lines 3-4 (high saponin) and 6-1 (low saponin), selected for detailed molecular analyses, are indicated.

Saponin content, assessed by a standard foaming assay, showed presence–absence variation: out of the 143 lines, 32 had detectable saponins, whereas 111 exhibited low to no foaming (Figure 1). Untargeted metabolomics via LC-qToF-MS/MS revealed 16 chemically distinct saponins derived from oleanolic, hederagenin, phytolaccagenic and serjanic acid (Table S1), considerably expanding the diversity previously reported from roots (three saponins; Lavaud et al., 2000) and aerial tissues (four saponins; Huong et al., 2021).

To represent the extremes of this variation, we selected two lines from the same field site (Taastrup) with similar protein content (∼20%) but contrasting saponin profiles. In lines 3-4 we detected nine saponins, whereas in line 6-1 we only detected four saponins albeit at lower levels (Table S2). Field trials of these two representative lines demonstrated promising agronomic performance: line 3-4 reached seed yields of up to 1.5 t/ha and line 6-1, 1.4 t/ha under Danish field conditions (Table S3), comparable to or exceeding reported seed yields for the related quinoa crop (*Chenopodium quinoa*) cultivated under North-West European field conditions, where varietal performance ranged from 0.47 to 3.42 t/ha (De Bock et al., 2021). The decoupling of protein and saponin content underscores the versatility of *C. album*: high-protein, low-saponin genotypes may be suitable for food or feed applications, while contrasting lines provide a valuable system to dissect the molecular basis of saponin biosynthesis and explore strategies to optimize it for different food and feed uses.

These findings provide the first comprehensive characterization of intraspecific variation in seed composition and specialized metabolite profiles in *C. album*, establishing it as a promising candidate for rapid domestication. By combining high nutritional potential with contrasting metabolite profiles, *C. album* offers a powerful system for both breeding and mechanistic studies of wild plant domestication. To explore the genetic basis of this phenotypic diversity, we assembled a reference-quality genome for *C. album* and examined ploidy structure across representative lines.

### A high-quality tetraploid genome assembly and ploidy variation in *C. album*

To establish a genomic reference for *C. album*, we generated a high-quality *de novo* genome assembly for the low-saponin tetraploid line 6-1 (Figure 2). Phylogenetic analysis placed *C. album* within the Amaranthaceae, closely related to *C. quinoa* (Figure 1A, Table S5), providing an evolutionary framework for comparative genomic and metabolic analyses. Using approximately 200 Gb of PacBio HiFi long reads (∼40× haploid coverage), we produced a large (1.63 Gb) and highly continuous genome assembly, consistent with GenomeScope estimates indicating extensive duplication expected for a tetraploid genome (Figure S1). The assembly captures nearly the full complement of expected plant genes, with BUSCO analysis identifying 2,239 of 2,326 conserved eudicot orthologs as complete, many present in duplicate (Figure 2). High contiguity is reflected by the small number of contigs (1,261) and large contig sizes (N50 50.4 Mb; N90 22.4 Mb), together with favorable QUAST metrics (GC content 36.2%; L50/L90 of 14/32) (Table S4). Together, these results establish a nearly complete, reference-quality genome assembly that provides a robust foundation for comparative and functional analyses.

**Figure 2.**
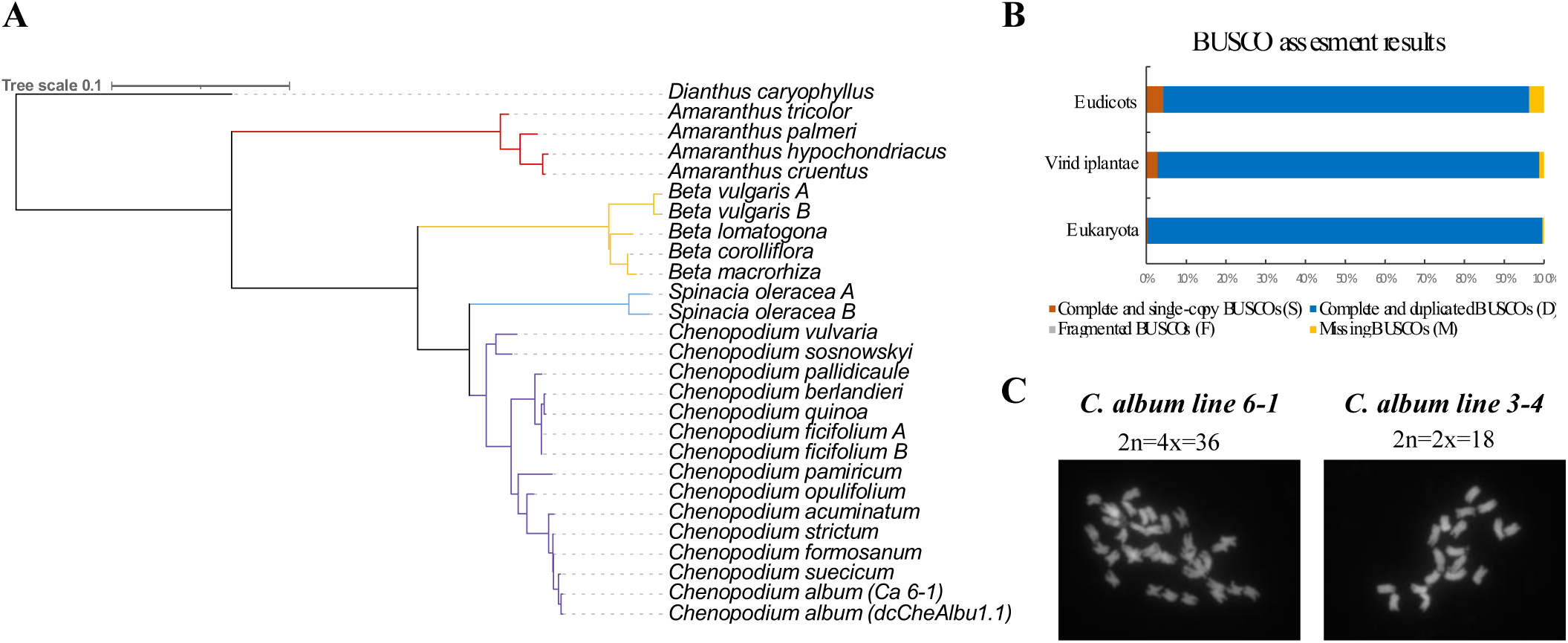
Genomic landscape of *Chenopodium album*. **A.** Phylogenetic placement of line 6-1 based on whole-genome comparison with representative Amaranthaceae species, with Dianthus (Caryophyllaceae) included as an outgroup. **B.** BUSCO completeness assessment of line 6-1 genome, showing a high proportion of complete and duplicated eudicot orthologs, consistent with a high-quality tetraploid assembly. **C.** Chromosome counts for the two focal accessions. Line 6-1 (low saponin) shows 36 (2n=4x=36) chromosomes, consistent with tetraploidy, whereas line 3-4 (high saponin) shows 18 (2n=2x=18) chromosomes, consistent with diploidy.

Because *C. album* is known for its complex evolutionary history and variable ploidy, we examined the genomic configuration of a subset of ten representative accessions subjected to ploidy analysis (Table S6). Flow cytometry and chromosome counts showed that line 6-1 is tetraploid (2n=4x=36; 3.5 ± 0.05 pg, 2C), whereas line 3-4 is diploid (2n=2x=18; 1.70 ± 0.07 pg, 2C) (Figure 2; Table S6). Among the remaining accessions analyzed for ploidy, only 95-5 was also diploid, while the others were tetraploid (Table S6; Figure S2). The two diploid lines analyzed exhibited high saponin and protein content, while no clear association was observed between ploidy level and saponin content across the remaining accessions. In this panel, ploidy alone therefore does not explain saponin variation, motivating further genomic analyses.

### Genome composition and sequence variation

To characterize the genomic landscape of *C. album*, we analyzed the composition and gene content of line 6-1. Transposable element annotation revealed that approximately 65% of the *C. album* genome consists of repetitive sequences, dominated by long terminal repeats (LTRs), and helitrons (Table S7). Evidence-based gene annotation integrating RNA-seq data identified 53,924 protein-coding genes of which 48,182 were functionally annotated with Gene Ontology (GO) and KEGG terms (Table S8; Files S1 and S2). The extensive gene complement and repeat fraction are consistent with the polyploid origin of *C. album* and provide a comprehensive foundation to investigate the genetic control of its metabolic diversity and other traits relevant to domestication.

To investigate sequence-level variation associated with saponin biosynthesis, we sequenced the high-saponin, diploid line 3-4 using the 6-1 genome as reference, achieving 98.9% read mapping at 90× coverage (Table S9, Figure S3). Variant calling identified 2.33 million single-nucleotide variants (SNVs) and small insertions or deletions (indels) across 23,829 genes, while structural variant (SV) analysis detected 63,270 SVs affecting 33,578 genes (Files S1 and S2). These results reveal extensive genomic variation, suggesting that saponin phenotypes are primarily driven by allelic and structural polymorphisms rather than ploidy level.

Collectively, the *C. album* line 6-1 reference genome provides a nearly complete, highly contiguous reference that captures the duplication and structural complexity characteristic of *C. album*. Together with ploidy and resequencing analyses, it establishes a comprehensive framework to dissect the genetic basis of nutritional and anti-nutritional traits. These genomic resources provide the basis for identifying key saponin biosynthetic genes, linking allelic variation to metabolite diversity, and guiding targeted domestication strategies in this resilient wild species.

### Identification and regulation of core saponin biosynthetic genes in *C. album*

To identify the genes underlying saponin accumulation in *C. album*, we integrated transcriptomics (Figure S4-6; Table S10-11), phylogenetic, and genomic analyses of the high-saponin diploid line 3-4 and the contrasting low-saponin tetraploid line 6-1. Comparative transcriptome profiling of leaves and seeds (Table S12) revealed generally higher gene expression in leaves than in seeds, and overall higher transcript abundance in the tetraploid 6-1 line (Figure S7-S8). Despite these global differences, GO enrichment analysis detected no significant functional category differences between tissues or lines, suggesting that saponin regulation involves a limited subset of genes.

Saponins are synthesized via the linear precursor 2,3-oxidosqualene, which is cyclized by oxidosqualene cyclases (OSCs) to generate a range of diverse triterpenoid scaffolds that are subsequently oxidized by cytochrome P450s and glycosylated by UDP-glycosyltransferases to yield bioactive saponins (Moses et al., 2014; Cárdenas et al., 2019). Homology searches against curated *C. quinoa* saponin biosynthetic genes (Jarvis et al., 2017) identified *C. album* orthologs encoding candidate OSCs, CYP716 cytochrome P450s, and glycosyltransferases in the 6-1 genome (Tables S13).

Phylogenetic and expression analyses identified β-amyrin synthases as the predominant OSCs in *C. album*, consistent with the dominance of oleanolic acid-derived saponins detected in metabolomic profiling (Table S1). These β-amyrin synthases showed strong seed-specific upregulation in the high-saponin line 3-4 relative to 6-1 (Figure 3; Table S14), supporting their central role in triterpenoid scaffold formation. Downstream oxidation steps are mediated by CYP716-family cytochromes P450, and nine putative orthologs were identified in the *C. album* genome that cluster with the CYP716A and CYP716B clades (Figure 3; Table S15). Members of CYP716s are known to catalyze C-28 oxidations required for oleanolic-acid aglycone formation, highlighting a direct correspondence between the predicted biosynthetic pathway and the saponin profiles observed in seeds.

**Figure 3.**
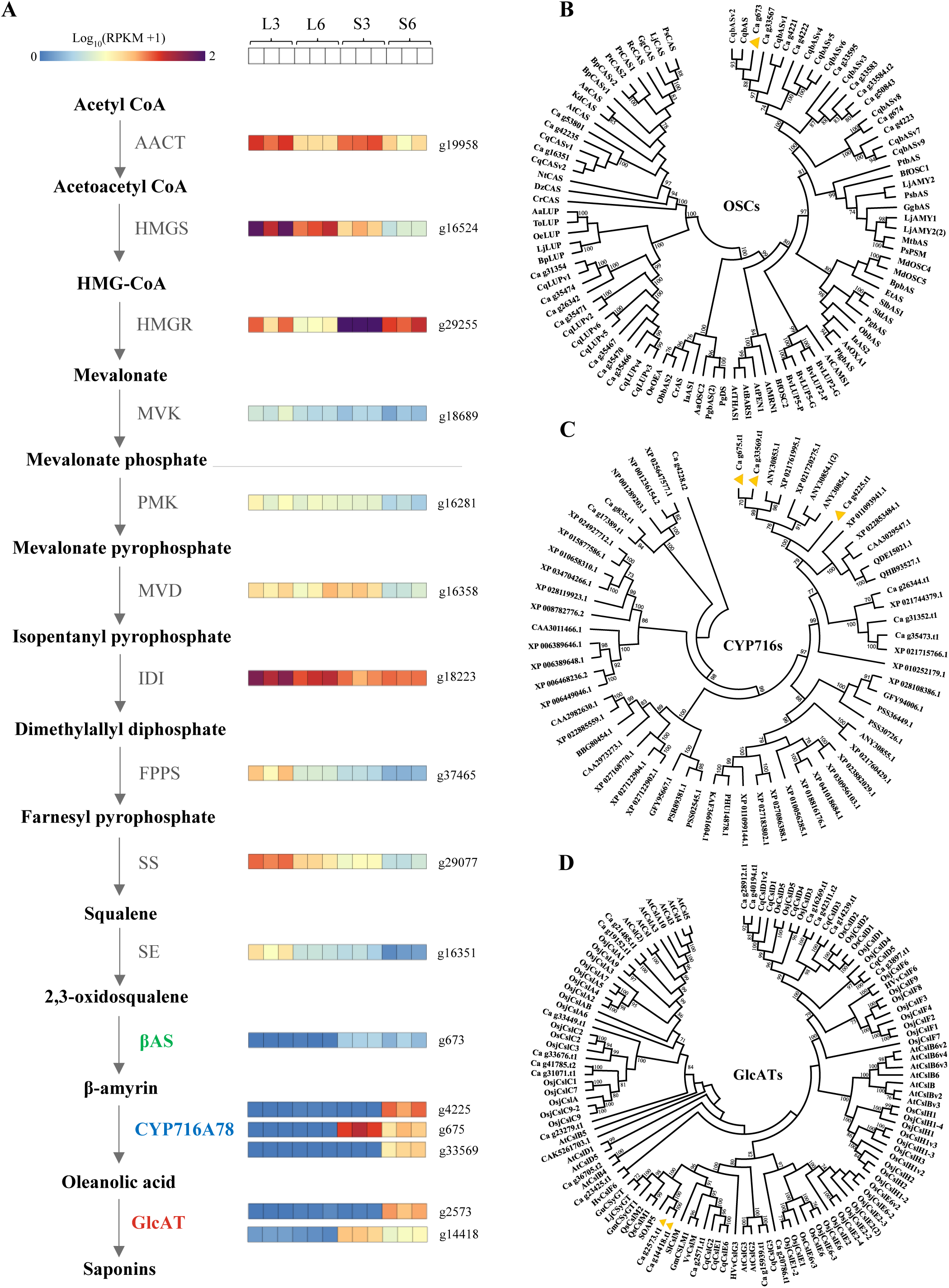
Expression and evolutionary context of the core saponin biosynthetic pathway in *Chenopodium album*. **A.** Simplified pathway and gene expression. Reconstructed biosynthetic route from 2,3-oxidosqualene to oleanolic acid-3-O-β-D-glucuronoside, showing only the predicted steps: (i) CaβAS (OSC) converts 2,3-oxidosqualene to β-amyrin; (ii) CYP716A78s oxidize β-amyrin to oleanolic acid; (iii) CaGlcATs catalyze glucuronidation to oleanolic acid-3-O-β-D-glucuronoside. Structures for the three key metabolites are shown. Expression heatmaps (log_10_[RPKM+1]) display transcript abundance for these genes in leaves (L) and seeds (S) of the high-saponin line 3-4 and low-saponin line 6-1 (three biological replicates per tissue and line). **B.** Phylogenetic placement of oxidosqualene cyclases (OSCs). Maximum-likelihood tree of OSC proteins from representative Amaranthaceae and related species. The validated *C. album* β-amyrin synthase (CaβAS) is highlighted. Bootstrap values (1,000 replicates) are shown for major nodes. **C.** Phylogeny of CYP716 family enzymes. Maximum-likelihood tree of CYP716 proteins across species. The validated *C. album* enzymes CYP716A78v1 and CYP716A78v2 are highlighted. Bootstrap support values (1,000 replicates) are indicated. **D.** Phylogeny of glucuronosyltransferases (GlcATs). Maximum-likelihood tree of CslM/GlcAT family members from representative Amaranthaceae species. CaGlcAT2, the enzyme responsible for seed saponin glucuronidation, is highlighted. Bootstrap values (1,000 replicates) are shown. The triangles in the figure represent genes from *C. album* functionally characterized in this study.

Finally, we examined glycosylation enzymes responsible for decorating triterpenoid aglycones. Glycosylation, a key structural diversification step in saponin biosynthesis, is catalyzed by UDP-glycosyltransferases (UGTs) and cellulose synthase-like M (CslM) proteins, which function as glucuronic-acid transferases (GlcATs). Sequence homology and phylogenetic analysis identified two *C. album* genes (g2573 and g14418) that cluster with known GlcATs involved in saponin glucuronidation (Figure 3; Table S15). Both genes are expressed in seeds, supporting a role in the biosynthesis of oleanolic-acid-3-O-glucuronides, the predominant saponins in line 3-4.

To assess sequence variation associated with differential saponin accumulation, we examined allelic diversity across the identified OSC, CYP716, and GlcAT genes, which uncovered 2,196 sequence variants, most of which were synonymous or non-coding. Only 11 missense variants with potential moderate effects were identified, associated with five genes: g24030.t1 (3-hydroxy-3-methylglutaryl coenzyme A reductase), g29077.t1 (squalene synthase), g673.t1 (terpene cyclase/mutase family member), g33583.t1 (terpene cyclase/mutase family member), and g34948.t2 (CYP716A; beta-amyrin 28-monooxygenase) (Table S16). Despite the lower ploidy of the high-saponin line 3-4, most saponin biosynthetic genes exhibited higher expression in its seeds relative to the low-saponin tetraploid 6-1 (Figure 3), a pattern inconsistent with a simple gene-dosage expectation and consistent with regulatory or sequence-specific differences contributing to saponin accumulation. Collectively, these findings define the core genetic architecture of the saponin biosynthetic pathway in *C. album*, integrating phylogenetic placement, tissue-specific expression, and allelic diversity. This framework identifies candidate genes for functional validation and provides a foundation for targeted metabolic engineering of saponins and the domestication of this resilient wild species.

### Functional validation of the core saponin biosynthetic pathway in *C. album*

In *C. album* seeds, oleanolic acid-derived saponins dominate the triterpenoid profile, with oleanolic acid monoglucuronoside, the simplest and most abundant product, representing the principal metabolite (Table S1). To elucidate the enzymatic basis of this pathway, we predicted and reconstituted the sequential reactions from β-amyrin to oleanolic acid-3-O-β-D-glucuronoside (Figure 4) using candidate enzymes identified from our genome and transcriptome analyses.

**Figure 4.**
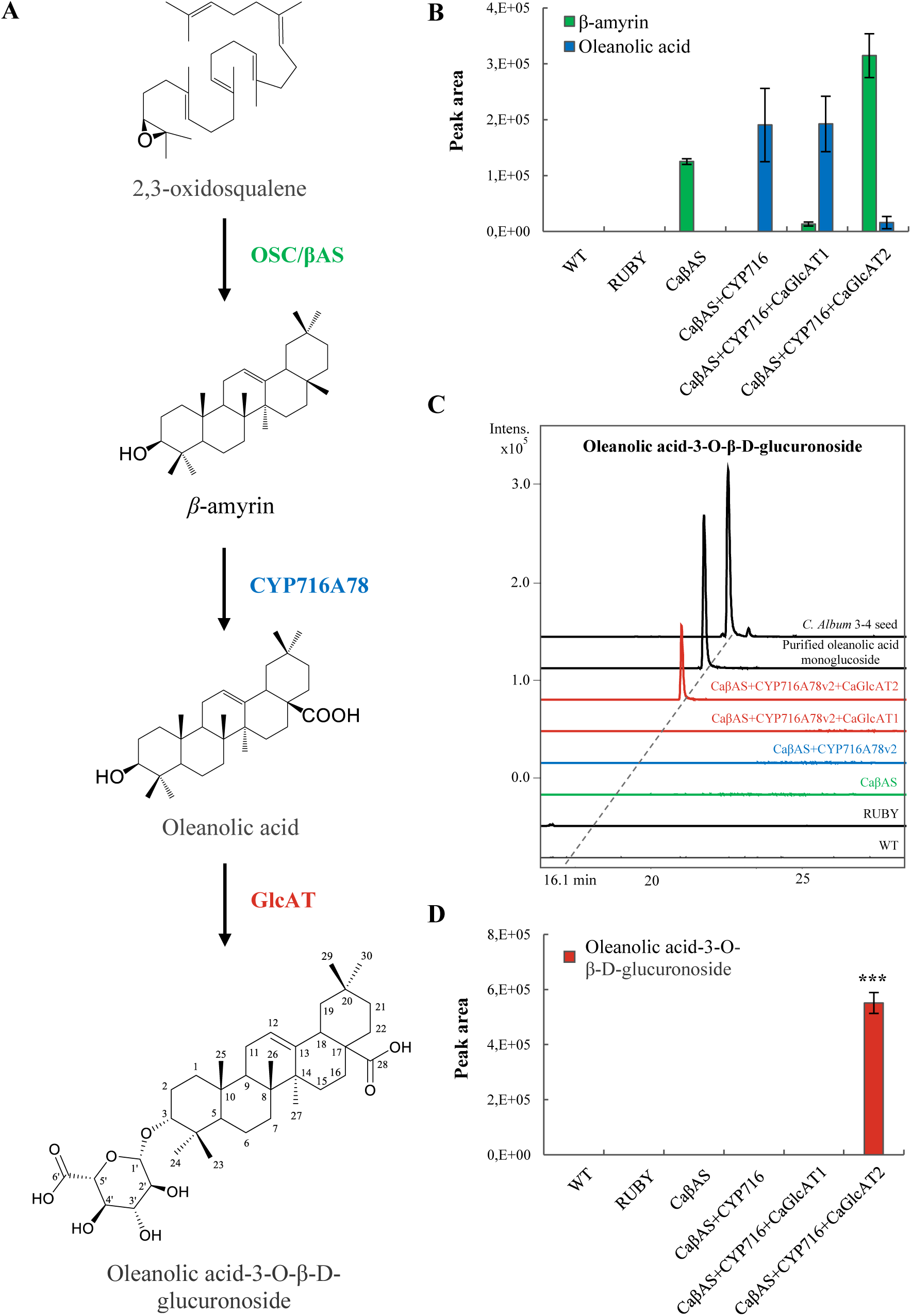
Functional reconstruction of the *Chenopodium album* saponin biosynthetic pathway in *Nicotiana benthamiana*. **A.** Enzymatic steps from 2,3-oxidosqualene to oleanolic acid-3-O-β-D-glucuronoside. Reconstitution of the *C. album* pathway in *N. benthamiana* using CaβAS (β-amyrin synthase), CYP716A78v2 (C-28 oxidase), and CaGlcAT2 (glucuronosyltransferase). The structure of oleanolic acid-3-O-β-D-glucuronoside was confirmed by 1D/2D NMR; selected COSY and HMBC correlations supporting the assignment are shown. **B.** GC–MS analysis of β-amyrin and oleanolic acid in infiltrated *N. benthamiana* leaves: wild type (WT), RUBY control, CaβAS, CaβAS + CYP716A78v2, CaβAS + CYP716A78v2 + CaGlcAT1, and CaβAS + CYP716A78v2 + CaGlcAT2. **C.** LC–MS chromatograms showing glucuronide production. Extracted-ion chromatograms for the same infiltration series demonstrating that oleanolic acid-3-O-β-D-glucuronoside accumulates only when CaGlcAT2 is co-expressed. **D.** LC–MS quantification of glucuronide formation. Peak-area quantification of the glucuronide product across the six treatments. All experiments were performed with three biological replicates.

A β-amyrin synthase (CaβAS; g673.t1) sharing 98.9% amino acid identity with the functionally characterized *C. quinoa* enzyme was strongly upregulated in seeds of the high-saponin line 3-4 compared with the low-saponin line 6-1 (Table S14). Phylogenetic analysis and its localization within a terpene biosynthetic gene cluster supported its role in triterpenoid scaffold formation (Figure 3; Figure S9; Table S17). Transient expression of *CaβAS* in *Nicotiana benthamiana* resulted in the production of β-amyrin, confirming its predicted enzymatic activity (Figure 4; Figure S10).

To characterize the oxidative steps from β-amyrin to oleanolic acid, we identified three *C. album* CYP716A homologs: g4225.t1 (CYP716A78v1), g675.t1 (CYP716A78v2), and g33569.t1 (CYP716A78v3), as candidate P450 enzymes. Co-expression of *CaβAS* with CYP716A78v1 or v2 transiently in *N. benthamiana* efficiently converted β-amyrin to oleanolic acid, whereas CYP716A78v3 produced only partial conversion (Figure 4; Figure S10). These results demonstrate that CaβAS, together with specific CYP716A enzymes, forms the core oxidation module for oleanolic acid biosynthesis in *C. album*.

The first glycosylation step converts oleanolic acid to oleanolic acid-3-O-β-D-glucuronoside ([M-H]^-^ 631.38 m/z corresponding to a calculated exact mass of 632.3924 for C_36_H_56_O_9_), which represents the major saponin accumulated in seeds of the high-saponin line 3-4. LC-qToF-MS/MS and 1D/2D NMR analyses and elemental composition (C_36_H_56_O_9_) confirmed its structure, consistent with previously reported oleanane-type glucuronides (Figure 4; Figure S11; Table S18). Two candidate glucuronosyltransferases, CaGlcAT1 and CaGlcAT2 (g2573, g14418), were identified based on sequence homology and phylogenetic clustering with known CslM-type GlcATs. Phylogenetic analysis placed CaGlcAT2 within the CslM subfamily (Figure 4), supporting its role in triterpenoid glucuronosylation. Variant analysis revealed one missense mutation in CaGlcAT2 with a predicted moderate functional effect, suggesting a role in saponin biosynthesis in *C. album* (Table S16). Functional validation in *N. benthamiana* showed that co-expression of CaβAS and CYP716A78v2 with CaGlcAT2, but not CaGlcAT1, resulted in production of oleanolic acid-3-O-β-D-glucuronoside (Figure 4). LC-MS confirmed glucuronide formation only in the presence of CaGlcAT2, while GC-MS showed simultaneous shifts in β-amyrin and oleanolic acid levels, consistent with increased flux toward glycosylated products (Figure 4). These results identify CaGlcAT2 as the enzyme catalyzing the first glycosylation step in seed saponin biosynthesis, consistent with its high expression in seeds of the high-saponin line (Figure 4; Figure S12).

In summary, these findings functionally validate the core saponin biosynthetic pathway in *C. album*, from β-amyrin formation to oleanolic acid-3-O-β-D-glucuronoside. The coordinated activities of CaβAS, CYP716A78v1/v2, and CaGlcAT2 reconstruct the biosynthetic route for oleanolic acid-type seed saponins, linking integrated metabolomic, transcriptomic, and genomic predictions with biochemical outcomes. This validation establishes a mechanistic framework for both metabolic engineering and fundamental studies of triterpenoid metabolism, while providing molecular targets to fine-tune saponin content and composition during the domestication of this resilient wild species.

Overall, this work provides the first high-quality genome for Danish *C. album* and a validated biosynthetic framework for seed triterpenoid saponins. The tetraploid 6-1 genome captures the extensive duplication and structural complexity of the genus, while comparative analyses with the high-saponin diploid line 3-4 reveal allelic and structural diversity underlying key nutritional and anti-nutritional traits. Functional reconstruction of the β-amyrin to oleanolic acid-3-O-β-D-glucuronoside pathway confirms the molecular basis of seed saponin biosynthesis and establishes a mechanistic foundation for trait optimization.

The combined genomic, metabolomic, and functional resources position *C. album* as a promising model for both fundamental and translational research on wild plant domestication. Danish accessions show high protein content (14–22%), variable saponin levels, and field yields exceeding one ton per hectare, demonstrating its nutritional and agronomic potential. Collectively, these resources open avenues for advanced breeding and bioengineering strategies, including allele mining, AI-assisted breeding, and targeted gene editing, to fine-tune agrifood traits and accelerate the domestication of *C. album* as a resilient, high-protein food crop for locally adapted agriculture.

Altogether, this work establishes *C. album* as a proof of concept that integrated omics and functional biology can transform a wild species into a crop within a single research framework, linking fundamental discovery to practical domestication.

## Methods

A brief overview is provided here. Detailed protocols, parameters, and references are available in supplemental information.

### Plant material and growth conditions

A wild *Chenopodium album* core collection was established from seeds collected across Denmark and advanced by single-seed descent to S4–S5. Plants were grown in a climate-controlled greenhouse (21 °C, 16 h light/8 h dark). Two *C. album* accessions differing in saponin content (3-4, high-saponin; 6-1, low-saponin) were selected for genome sequencing, transcriptomics, metabolomics, protein and saponin quantification. Leaves and mature seeds were collected from S4 plants, with three biological replicates per tissue and line.

### Saponin and protein quantification

Seed saponin content was estimated using a standard foam test (Koziol, 1991). Protein content was calculated from total nitrogen measured by CHNS elemental analysis using a nitrogen-to-protein conversion factor of 6.25.

### Metabolite extraction and LC–MS

Leaf and seed samples were homogenized in liquid nitrogen and extracted in 85% methanol. Extracts were centrifuged and filtered prior to LC–MS analysis. Chromatographic separation used a reversed-phase C18 gradient and high-resolution qToF mass spectrometry in negative ion mode, following a method adapted from Khakimov et al. (2015). Data were processed with Bruker DataAnalysis.

### Ploidy, chromosome counts, and genome size

Ploidy levels and genome size were determined by flow cytometry using DAPI and propidium iodide staining with internal standards (Greilhuber, 2005; Van Oost et al., 2021). Chromosome counts were performed on colchicine-treated root tips using the Steamdrop method for high quality plant chromosome preparations (Kirov et al., 2014) and DAPI staining, followed by fluorescence microscopy and chromosome number estimation using DRAWID software.

### Genome sequencing, assembly, and annotation

High-molecular-weight DNA was extracted from mature leaves of the low-saponin tetraploid line 6-1, and a PacBio HiFi long-read whole-genome shotgun library was prepared and sequenced on the Sequel II platform. *De novo* assembly was generated using Hifiasm, polished with Racon, and screened for contaminants using Kraken2. GenomeScope2 was used to estimate genome size and ploidy from k-mer profiles. The assembled genome was soft-masked using EDTA for transposable-element annotation. Gene prediction was performed with BRAKER3, integrating RNA-seq data and Viridiplantae proteins, and functional annotation was carried out with PANNZER2.

### Variant calling and structural variation

Genomic DNA for Illumina short-read sequencing was extracted from leaves of the high-saponin diploid line 3-4. Reads were trimmed and mapped to the *C. album* 6-1 reference genome using minimap2. Variants were called with Platypus and filtered with bcftools. Structural variants were identified with Manta using default and size-based thresholds. Variant effects were annotated with VEP using the *C. album* genome annotation.

### RNA-seq and differential expression analysis

Total RNA was extracted from leaves and seeds of lines 3-4 and 6-1 (three biological replicates per tissue and line). Twelve RNA-seq libraries were sequenced on the Illumina NovaSeq platform. Reads were trimmed with Trimmomatic and quantified using kallisto. Differential expression analysis was performed with DESeq2 and gene ontology enrichment with clusterProfiler.

### Identification of saponin biosynthetic genes in *C. album*

Genes involved in triterpenoid saponin biosynthesis in *C. album* were identified using homology-based searches in the *C. album* line 6-1 proteome against a curated dataset of protein sequences of known OSCs, CYP716s, and GlcATs from different species.

### Phylogenetic and cluster analysis

Species-level phylogenetic inference was performed using BUSCO genes as input for ASTRAL-Pro3 (Zhang et al., 2020), based on genome assemblies listed in Table S5. For gene family phylogenetic analyses, protein sequences were curated from published datasets obtained from NCBI or UniProt (Table S15). Maximum-likelihood trees were constructed using MEGA with 1,000 bootstrap replicates (Kumar et al., 2024). Biosynthetic gene clusters were predicted using plantiSMASH.

### Functional validation in *Nicotiana benthamiana*

Candidate OSC, CYP716, and GlcAT genes were cloned into pEAQ-HT vectors (primers listed in Table S19; cloned gene sequences in Data S1) and transiently expressed in *N. benthamiana* using *Agrobacterium tumefaciens* AGL1. Metabolites from infiltrated leaves were analyzed by GC–MS and LC–MS.

## Data availability

The genome assembly and annotation of *Chenopodium album* line 6-1 have been deposited in the NCBI under BioProject PRJNA1377043. Raw PacBio HiFi genome sequencing reads, Illumina resequencing data, and RNA-seq data have been deposited in the NCBI Sequence Read Archive and Gene Expression Omnibus under the same BioProject. Metabolomics data supporting this study are provided in the Supplementary Information. Additional data are available from the corresponding author upon reasonable request.

## Author contributions

S.V. and P.D.C. conceived the study and research plans. S.V. led the genome assembly, transcriptomic analyses, variant calling, metabolomics and multi-omics integration. A.S.M. led the field trial and contributed to data analysis, figure development, cross-validation of results, and manuscript revision. A.S.M. together with S.V., K.E. and B.C.C. performed gene cloning and functional characterization in *N. benthamiana*. J.G. and M.G.R. carried out saponin purification and NMR structural elucidation. K.V.L. and L.L. conducted and supported ploidy determination and chromosome counting. P.E.J. and S.B. contributed to the design of protein and saponin analyses, respectively. S.V. and P.D.C. wrote the manuscript with input from all authors. P.D.C. coordinated and supervised the project.

## Acknowledgments

We thank Thure P. Hauser for help in the initial identification of *C. album* in Denmark. We are grateful to Dorte Bodin Dresbøll for guidance and support during the field experiments, and we thank the greenhouse and field staff at the University of Copenhagen for their assistance with plant cultivation and maintenance. We also thank David Neilson for the CYP716 naming used in this study. NMR data were recorded at cOpenNMR, an infrastructure supported by the Novo Nordisk Foundation (NNF18OC0032996).

## Declaration of generative AI and AI-assisted technologies in the writing process

During the preparation of this work the authors used ChatGPT in order to improve readability of the manuscript. After using this tool, the authors reviewed and edited the content as needed and take full responsibility for the content of the publication.

