## Supplemental Data for "A genomic and functional framework for the rapid domestication of the wild plant *Chenopodium album*"

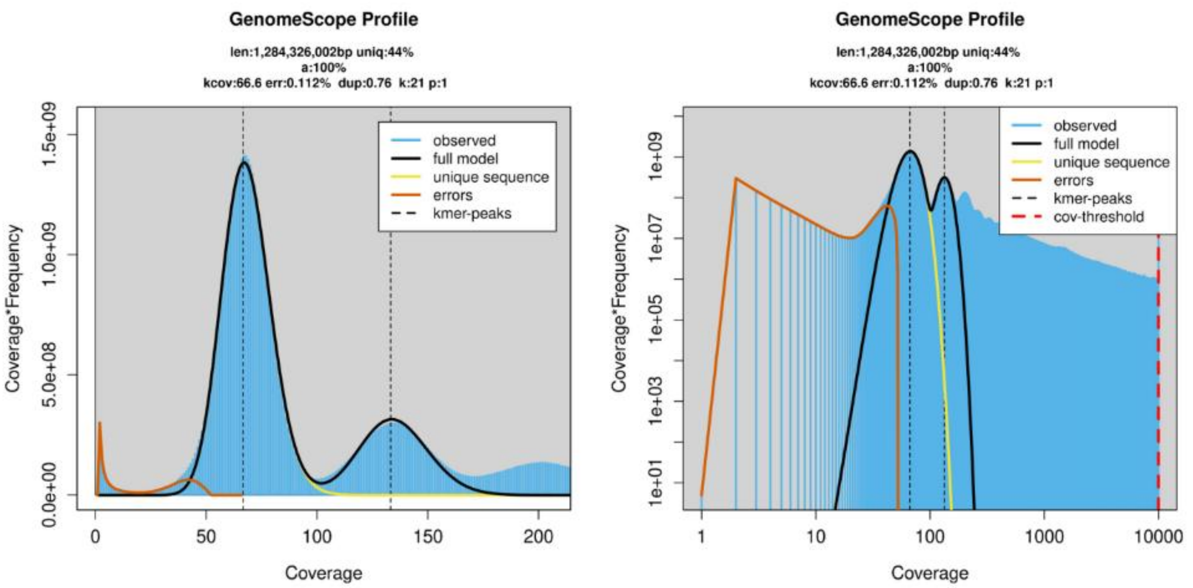


Results

```
GenomeScope version 2.0
input file = user_uploads/JPrqjre5KQ7TZ6Z8u4YG
output directory = user_data/JPrqjre5KQ7TZ6Z8u4YG
p = 1
k = 21

property                  min                 max
Homozygous (a)            100%                100%
Genome Haploid Length     1,279,979,090 bp    1,284,326,002 bp
Genome Repeat Length      716,762,589 bp      719,196,773 bp
Genome Unique Length      563,216,501 bp      565,129,230 bp
Model Fit                 54.3792%            90.0439%
Read Error Rate           0.111853%           0.111853%
```

Model

```
Formula: y_transform ~ x^transform_exp * length * predict1_0(k, d, kmercov,
    bias, x)

Parameters:
        Estimate Std. Error t value Pr(>|t|)
d       1.386e-01  1.779e-03   77.92   <2e-16 ***
kmercov 6.662e+01  5.647e-02 1179.83   <2e-16 ***
bias    7.599e-01  1.828e-02   41.56   <2e-16 ***
length  6.549e+08  3.113e+06  210.39   <2e-16 ***
---
Signif. codes:  0 '***' 0.001 '**' 0.01 '*' 0.05 '.' 0.1 ' ' 1

Residual standard error: 23590000 on 1996 degrees of freedom

Number of iterations to convergence: 6
Achieved convergence tolerance: 1.49e-08
```

**Figure S1. GenomeScope analysis of the *Chenopodium album* line 6-1 genome using Illumina short-read data.**

GenomeScope analysis was performed on Illumina paired-end reads from *C. album* line 6-1 using a k-mer size of 21. K-mer frequency distributions are shown on linear (left) and logarithmic (right) scales. The profiles display peaks corresponding to different k-mer coverage classes, enabling approximate inference of genome characteristics. The model estimated a haploid genome size of ~1.28 Gb, a mean k-mer coverage of ~66.6×, and a low sequencing error rate (~0.11%). Approximately 44% of the genome was inferred to consist of unique sequence, with a substantial proportion of duplicated content. Model parameters and summary statistics are shown below the plots. Given the duplicated and polyploid nature of the *C. album* genome, GenomeScope estimates should be interpreted as approximate.

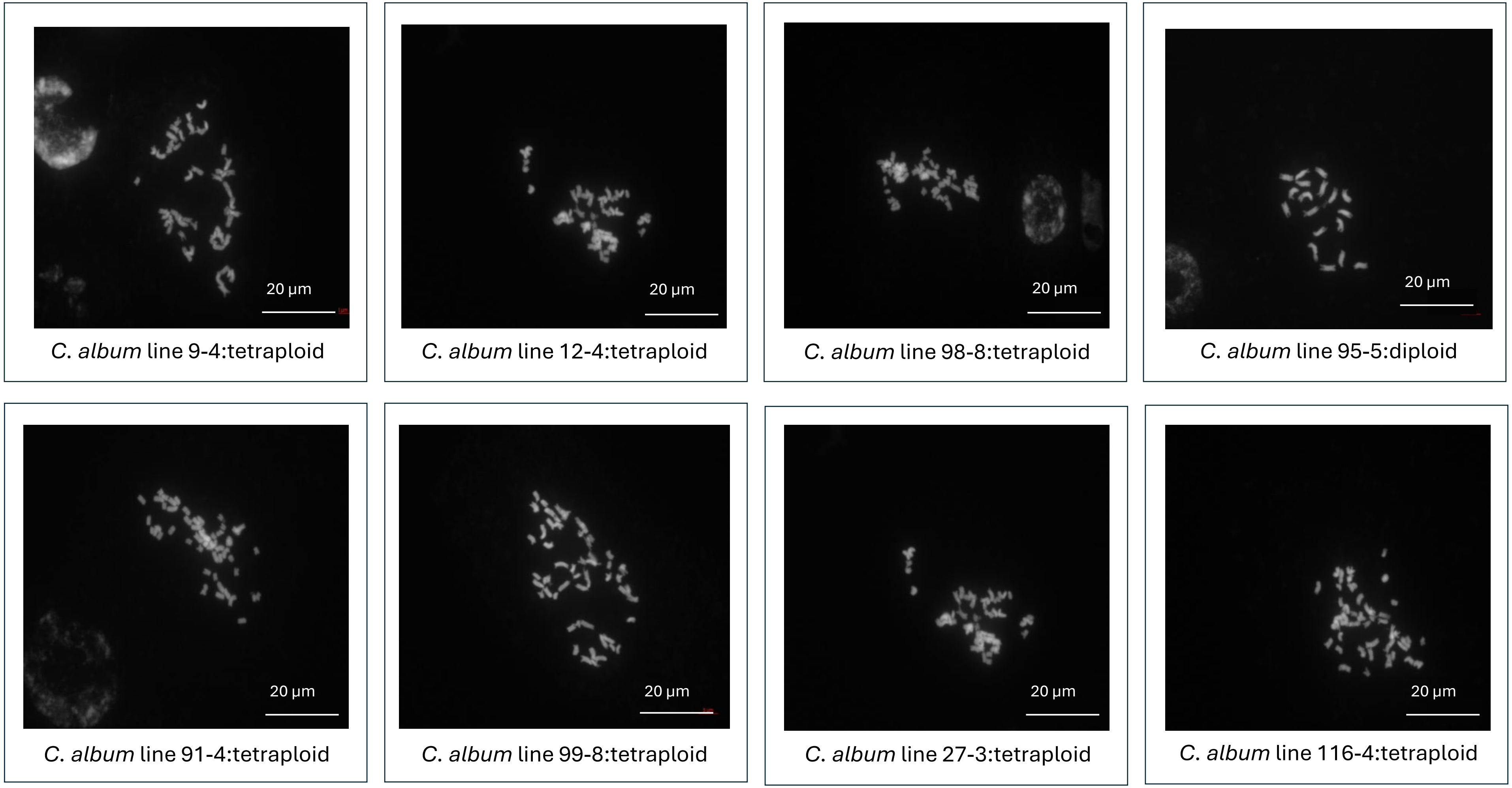


**Figure S2. Chromosome counts and ploidy variation among *C. album* lines.**

Representative DAPI-stained metaphase chromosome spreads from eight *C. album* lines collected across Denmark are shown. The analyzed lines include 12-4 and 9-4 from Taastrup; 98-8, 95-5, 91-4, and 99-8 from Stenlille; 27-3 from Rønne; and 116-4 from Silkeborg. Chromosome counts revealed ploidy levels ranging from diploid (2n = 2x = 18) to tetraploid (2n = 4x = 36) among the examined lines, demonstrating substantial intraspecific variation in chromosome number and genome organization.

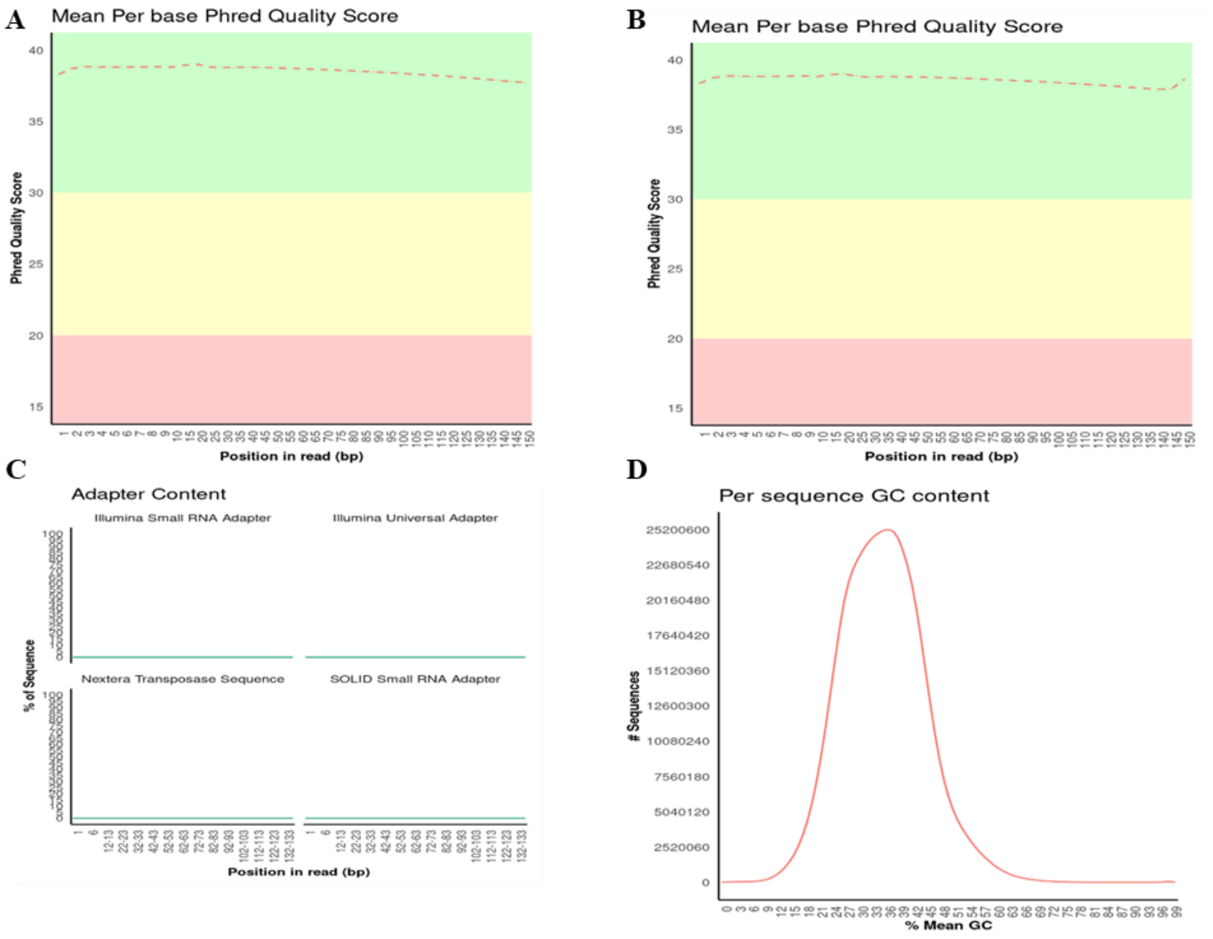


**Figure S3. Quality assessment of Illumina sequencing for *C. album* line 3-4.**

**A, B.** Mean per-base Phred quality scores for read 1 and read 2 across the paired-end Illumina dataset. Quality scores remain high across the read length, with only a modest decline toward the 3′ ends.

**C.** Adapter content across read positions. No substantial adapter-derived signal is detected across the read length.

**D.** Per-sequence GC-content distribution for the dataset, showing a single unimodal distribution.

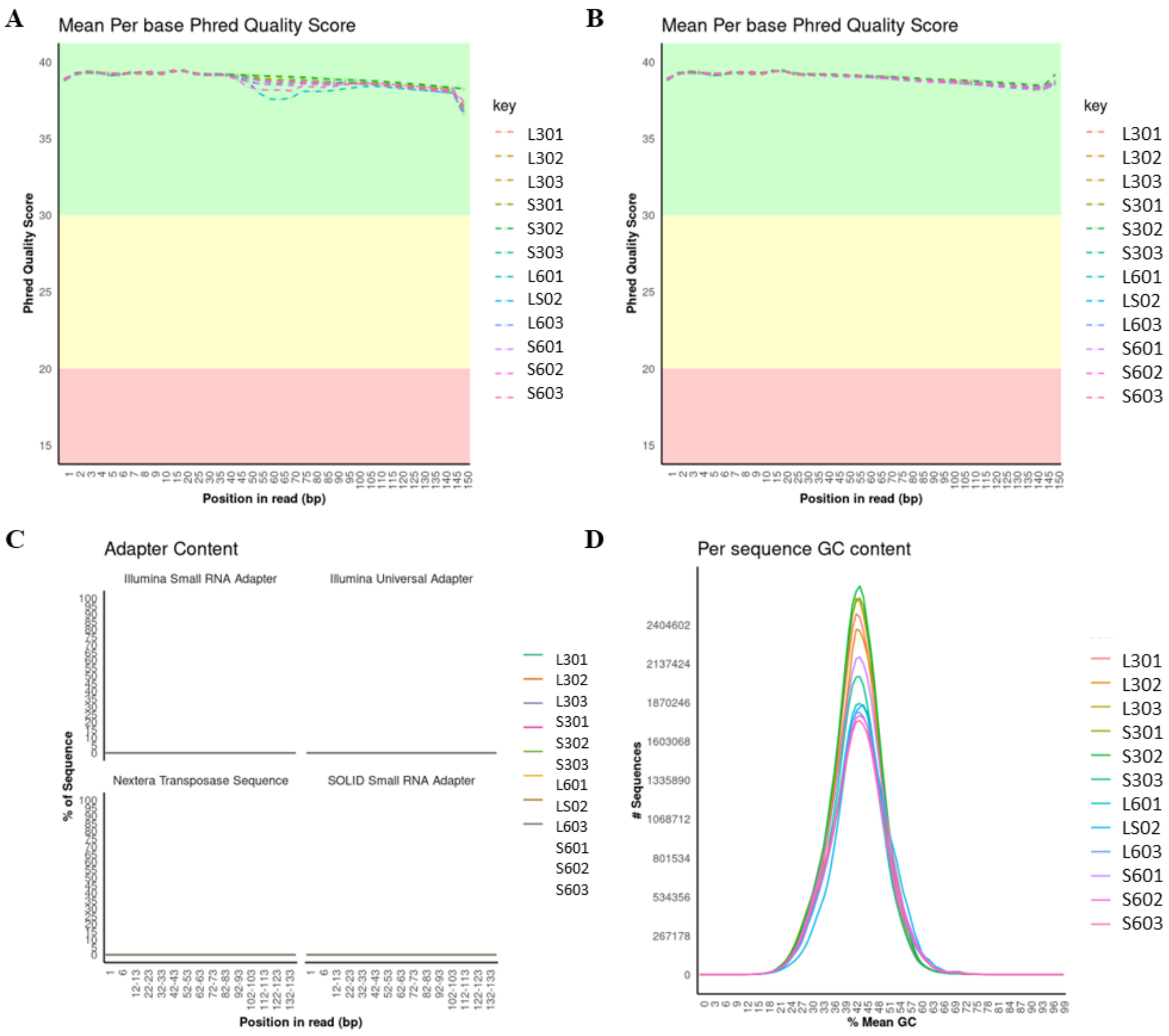


**Figure S4. Illumina sequencing quality assessment for transcriptomic samples of *C. album*.**

**A, B.** Mean per-base Phred quality scores across all transcriptomic libraries for read 1 and read 2. Quality scores remain high across the read length, with a modest decline toward the 3′ ends.

**C.** Adapter content across read positions for all libraries, showing no substantial adapter-derived signal.

**D.** Per-sequence GC-content distributions for all samples, displaying unimodal profiles with similar shapes across libraries.

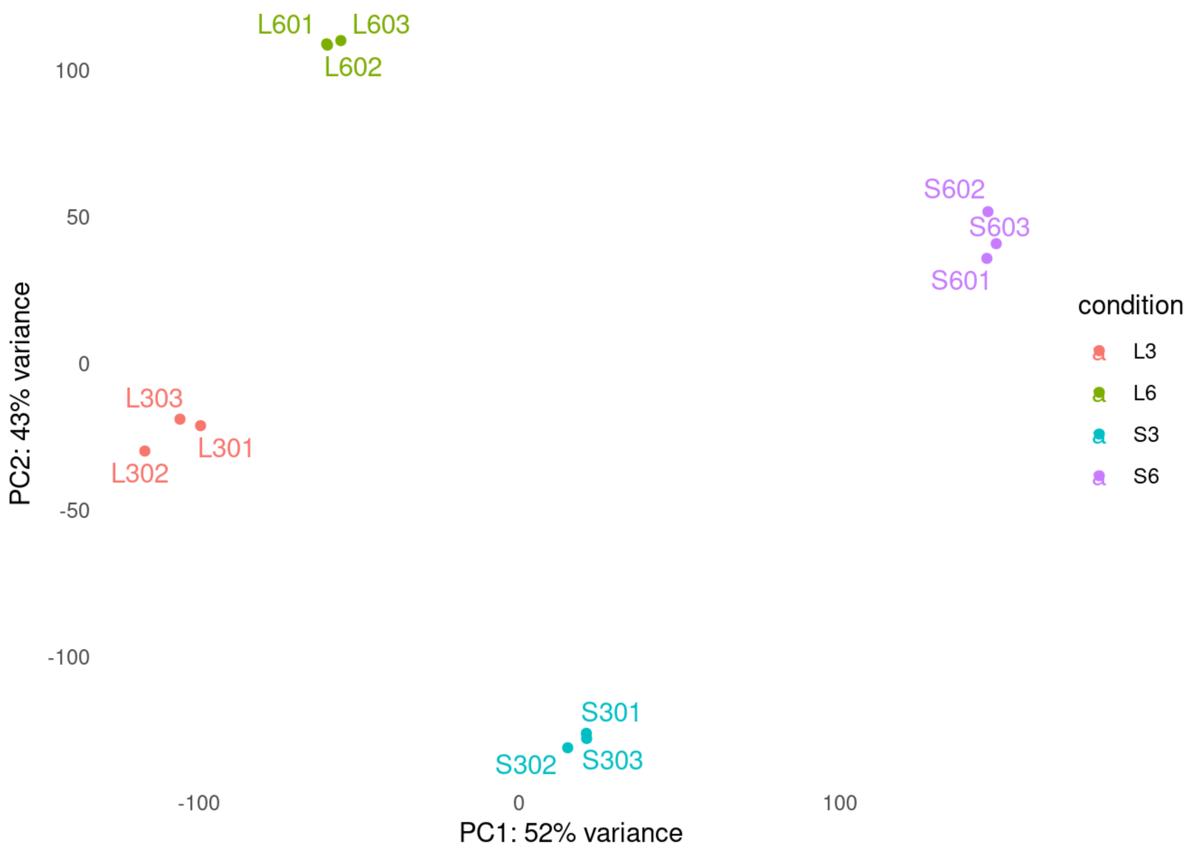


**Figure S5. Principal component analysis (PCA) of transcriptomic data from *C. album*.** Principal component analysis (PCA) of Illumina RNA-seq data from leaf and seed tissues of *C. album* lines 3-4 and 6-1 is shown. Each point represents one biological replicate. Samples are grouped by tissue and line: leaves from line 3-4 (L3; L301–L303), leaves from line 6-1 (L6; L601–L603), seeds from line 3-4 (S3; S301–S303), and seeds from line 6-1 (S6; S601–S603). The first principal component (PC1) explains 52% of the variance and the second principal component (PC2) explains 43% of the variance.

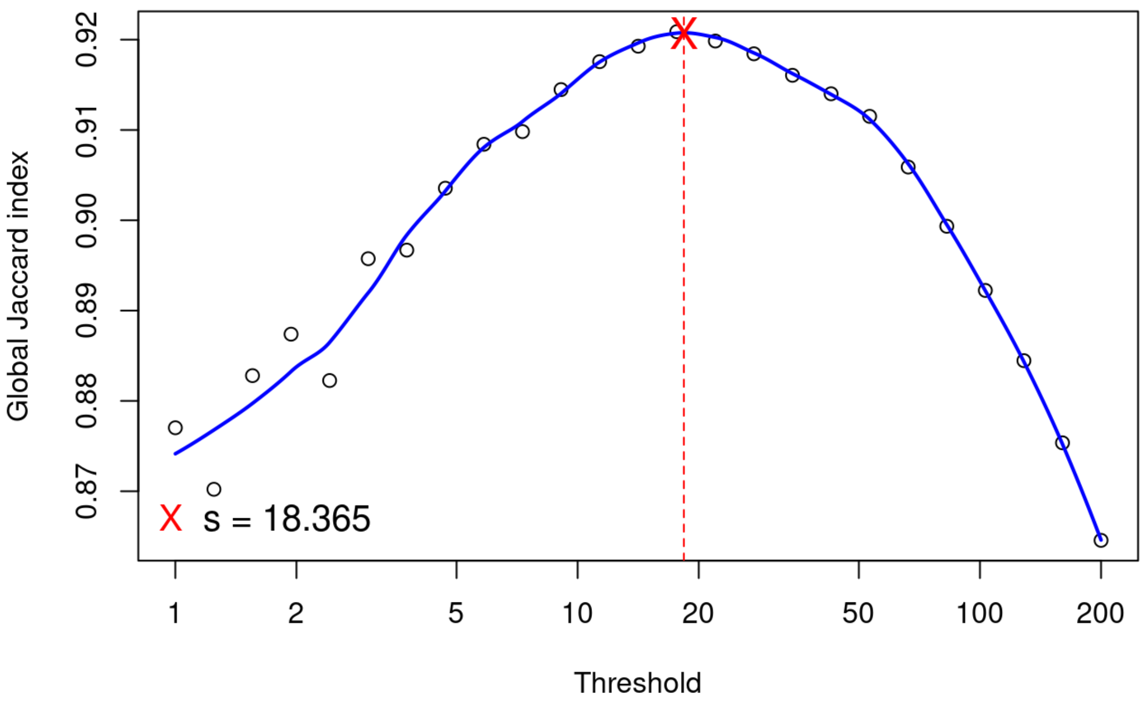


**Figure S6. Jaccard index of Illumina short-read data from *C. album* RNA-seq.**

The plot shows the Jaccard similarity index calculated from Illumina short-read datasets used in the transcriptomic analysis. The blue curve represents the Jaccard index across the evaluated range. The maximum Jaccard index is indicated by a red marker, with the corresponding position highlighted by a vertical dashed line.

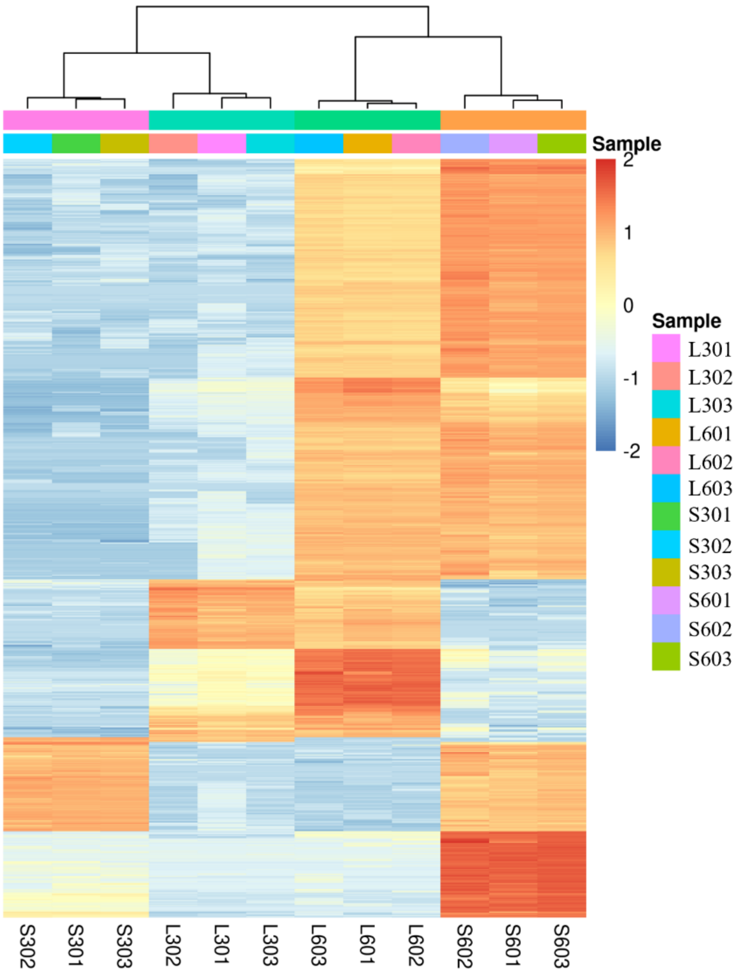


**Figure S7. Heatmap of the 1,000 most variable genes based on variance-stabilized RNA-seq data from *C. album*.**

Variance-stabilizing transformation (VST) was applied to Illumina RNA-seq count data, and the 1,000 genes with the highest variance across all samples were selected for visualization. Expression values for each gene were centered across samples. The heatmap displays hierarchical clustering of genes and samples based on these centered expression values. Sample identities (L301–L303, L601–L603, S301–S303, S601–S603) are indicated by color-coded bars. Warmer colors represent higher centered expression values, whereas cooler colors represent lower values.

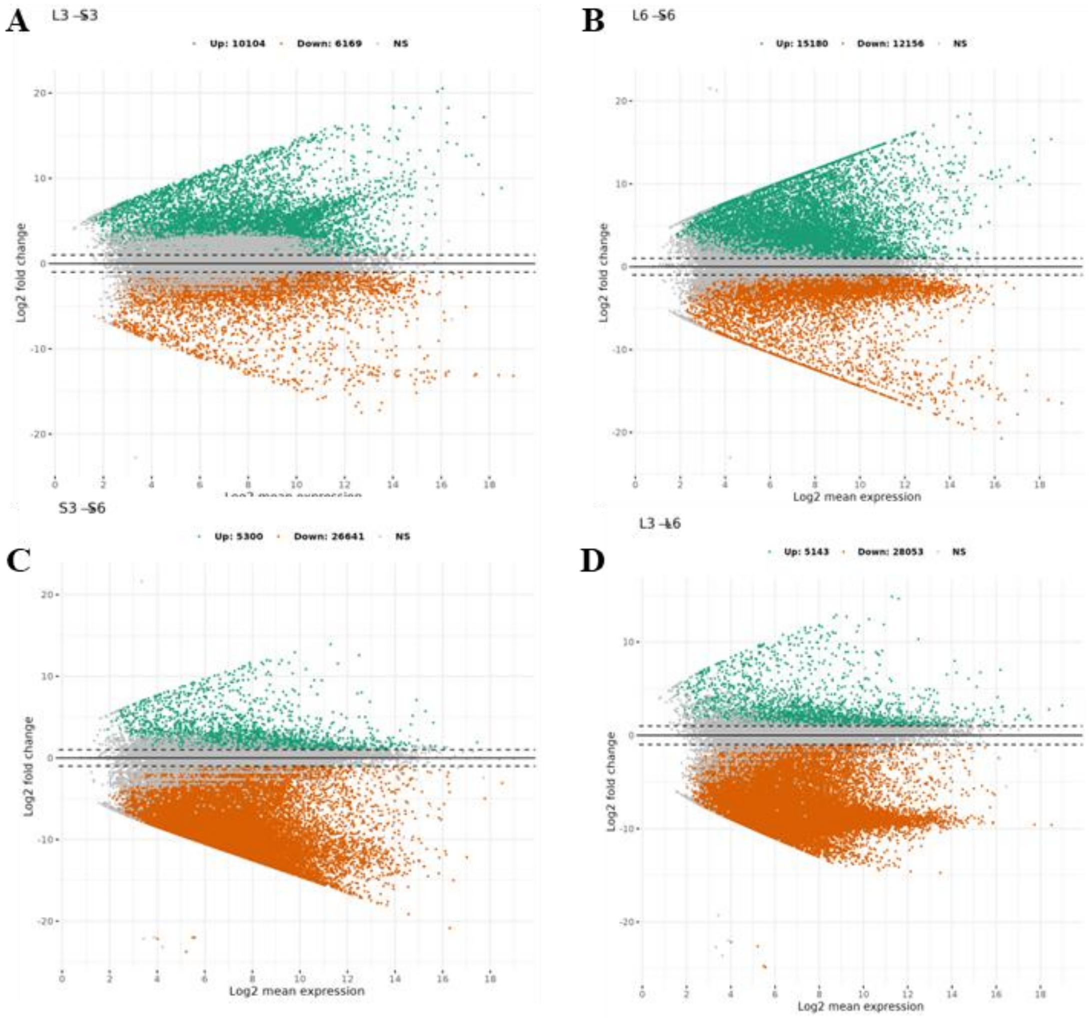


**Figure S8. MA plots of differential expression analyses in *C. album*.**

MA plots showing log2 fold change versus log2 mean expression for pairwise comparisons between tissues and lines.

**A.** Leaves versus seeds of *C. album* line 3-4 (L3 vs S3).

**B.** Leaves versus seeds of *C. album* line 6-1 (L6 vs S6).

**C.** Seeds of line 3-4 versus seeds of line 6-1 (S3 vs S6).

**D.** Leaves of line 3-4 versus leaves of line 6-1 (L3 vs L6).

Each point represents one gene. Genes identified as differentially expressed according to the applied statistical thresholds are highlighted, while non-significant genes are shown in grey. Dashed horizontal lines indicate the fold-change thresholds used in the analysis.

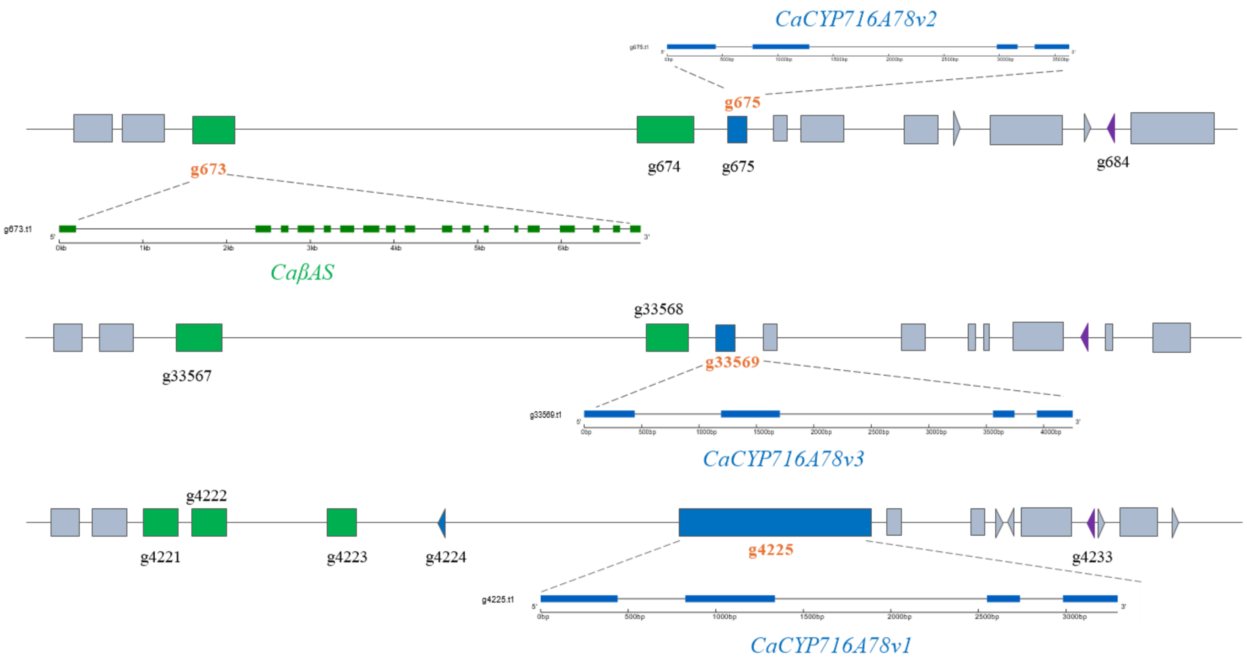


**Figure S9. Terpene biosynthetic gene clusters in *C. album*.**

Genomic organization of terpene biosynthetic gene clusters identified in the *C. album* line 6-1 genome using plantiSMASH (Kautsar et al., 2017). The positions of the functionally validated genes β-amyrin synthase (CaβAS; g673.t1) and cytochrome P450 enzymes of the CYP716A78 family (CaCYP716A78v1, g4225.t1; CaCYP716A78v2, g675.t1; CaCYP716A78v3, g33569.t1) are shown within their respective genomic clusters. Gene models are displayed with coding regions indicated, and neighboring genes within each cluster are shown to illustrate local genomic context. Green boxes correspond to oxidosqualene cyclases, blue boxes to cytochrome P450s associated with terpene metabolism, and purple boxes to prenyltransferases (see details in Table S17).

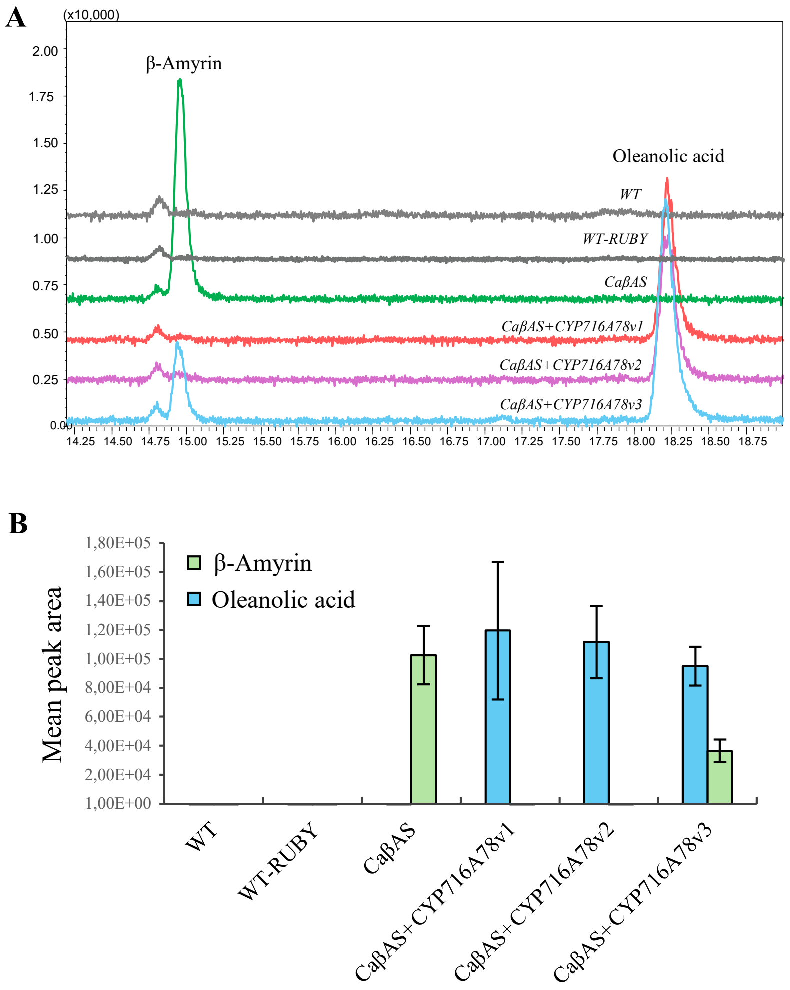


**Figure S10. GC–MS analysis of β-amyrin and oleanolic acid production following heterologous expression of *C. album* genes in *Nicotiana benthamiana*.**

**A.** Overlay of GC–MS chromatograms from *N. benthamiana* leaves transiently expressing saponin biosynthetic genes from *C. album*. Traces correspond to wild type (WT), β-amyrin synthase alone (CaβAS), and CaβAS co-expressed with individual CYP716A78 enzymes (CaβAS+CYP716A78v1, CaβAS+CYP716A78v2, CaβAS+CYP716A78v3). Peaks corresponding to β-amyrin and oleanolic acid are indicated.

**B.** Quantification of GC–MS peak areas for β-amyrin and oleanolic acid across the indicated *N. benthamiana* samples. Bars represent mean peak area ± s.e. (n=3).

**A**

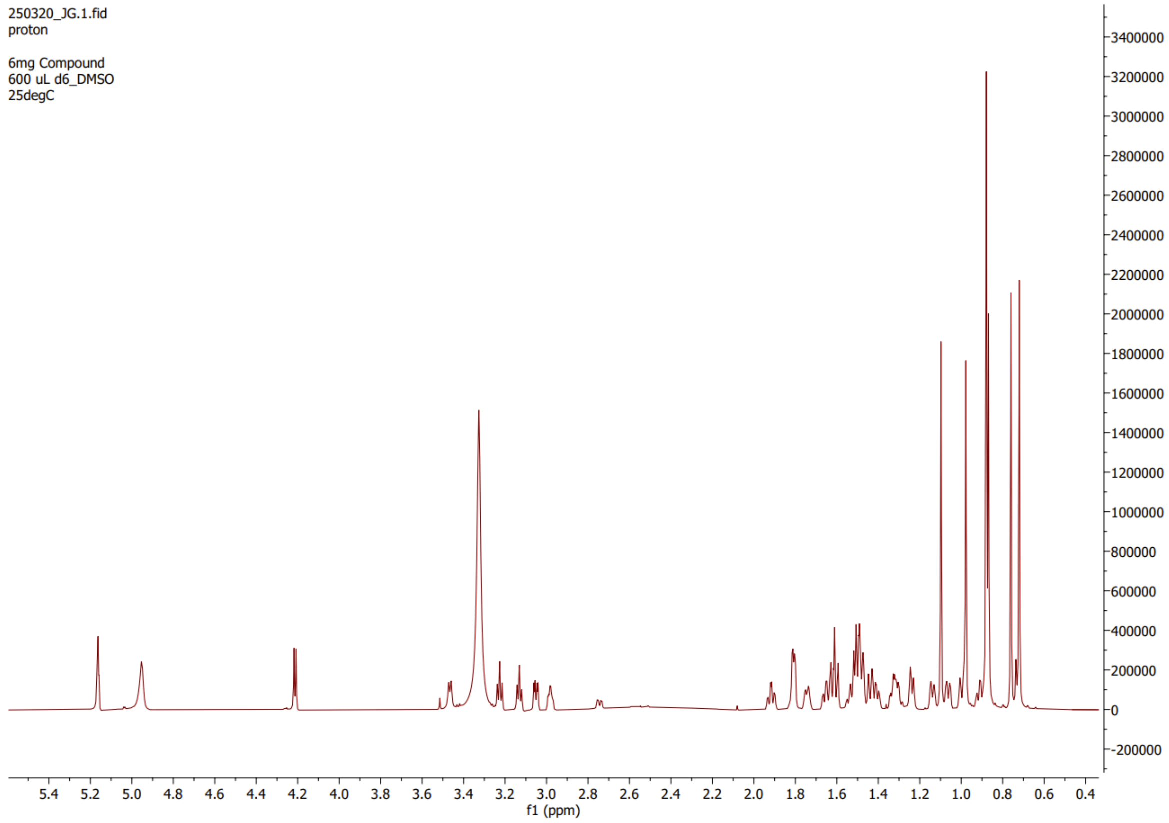

250320_JG.1.fid
proton

6mg Compound
600 uL d6_DMSO
25degC
3400000
3200000
3000000
2800000
2600000
2400000
2200000
2000000
1800000
1600000
1400000
1200000
1000000
800000
600000
400000
200000
0
-200000
5.4 5.2 5.0 4.8 4.6 4.4 4.2 4.0 3.8 3.6 3.4 3.2 3.0 2.8 2.6 2.4 2.2 2.0 1.8 1.6 1.4 1.2 1.0 0.8 0.6 0.4
f1 (ppm)


**B**

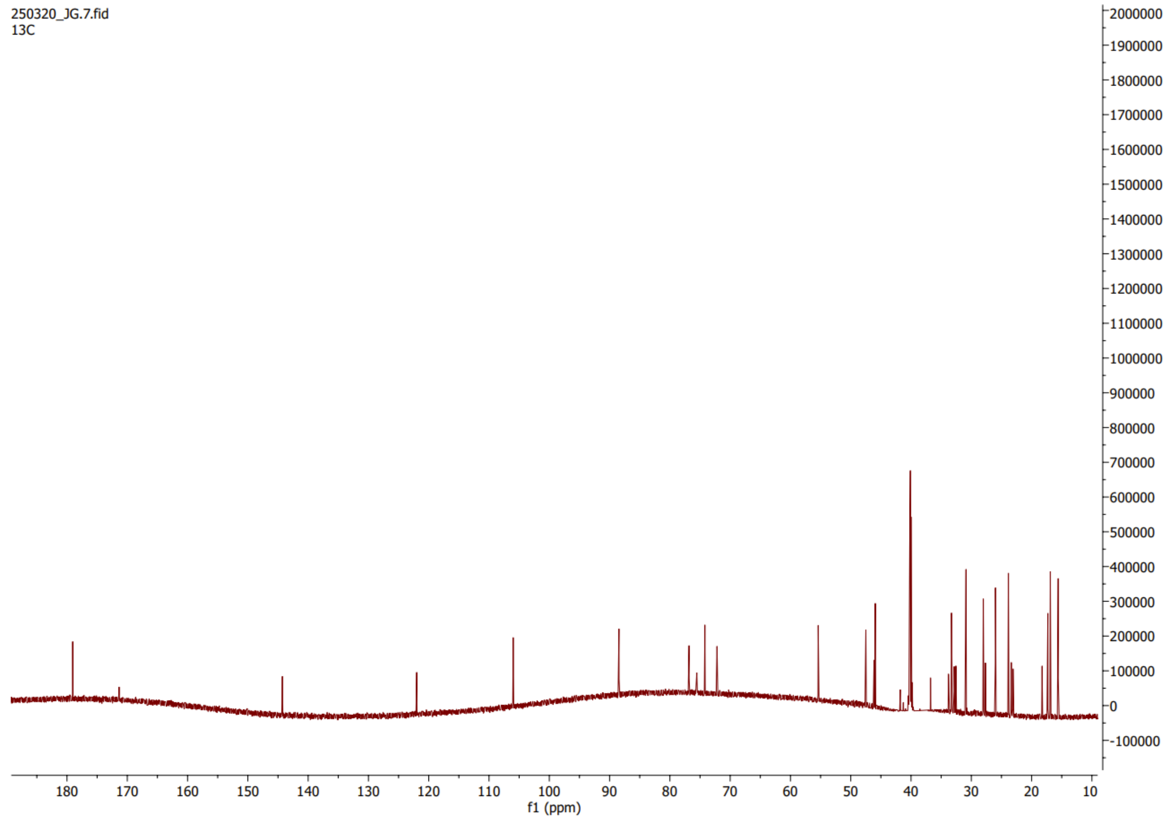

250320_JG.7.fid
13C
2000000
1900000
1800000
1700000
1600000
1500000
1400000
1300000
1200000
1100000
1000000
900000
800000
700000
600000
500000
400000
300000
200000
100000
0
-100000
180 170 160 150 140 130 120 110 100 90 80 70 60 50 40 30 20 10
f1 (ppm)

**C**

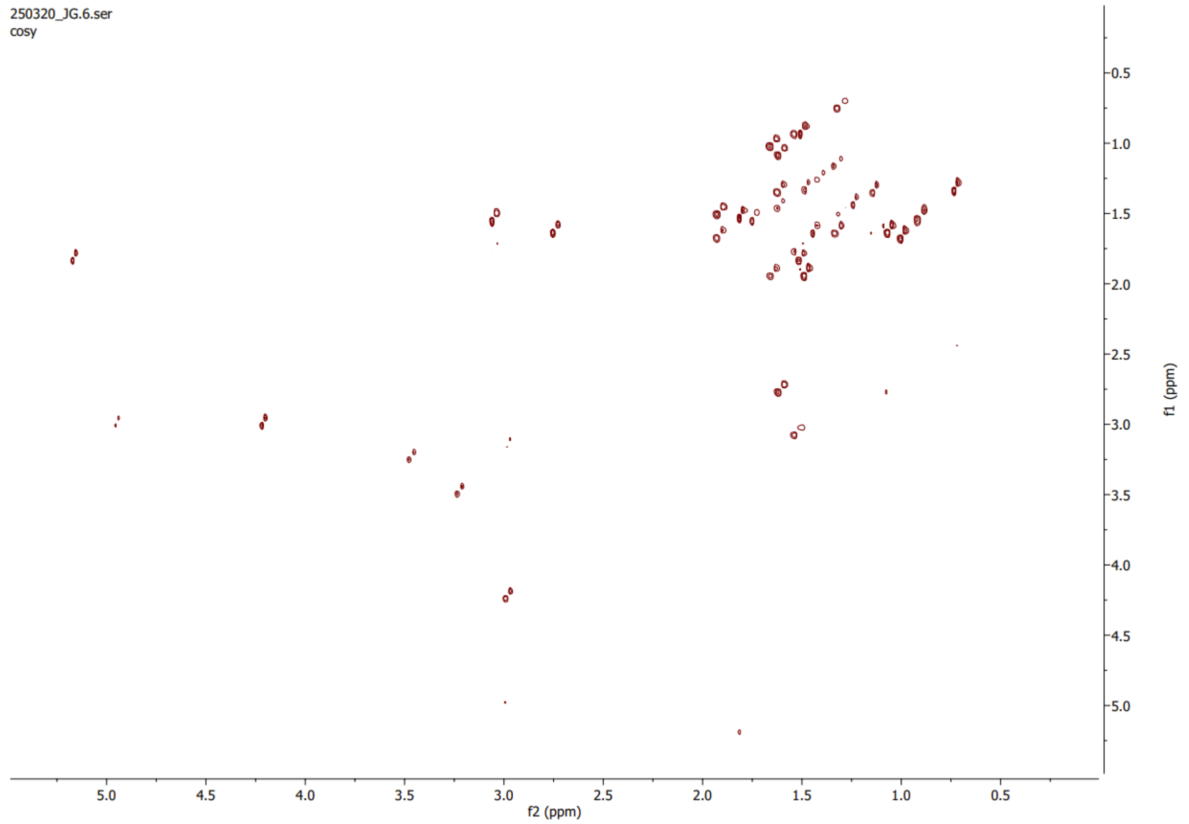

250320_JG.6.ser
cosy
f2 (ppm)
f1 (ppm)


**D**

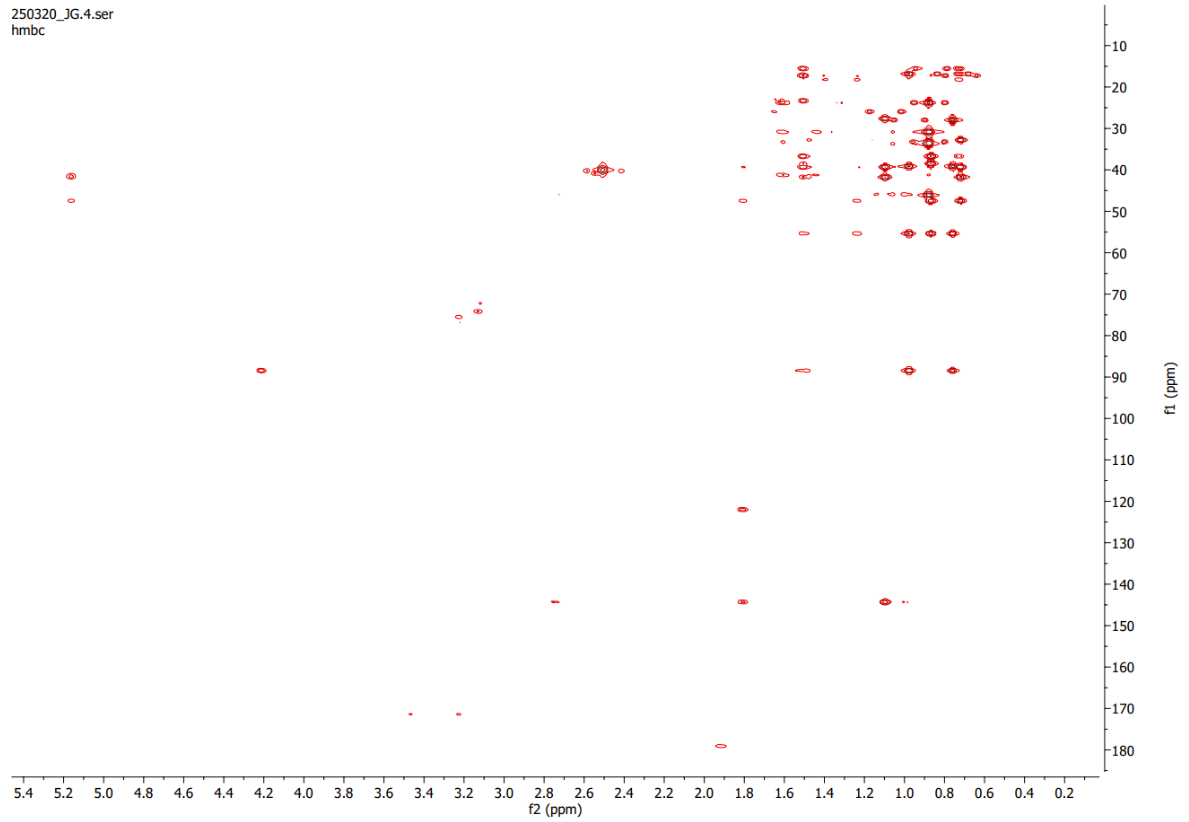

250320_JG.4.ser
hmbc
f2 (ppm)
f1 (ppm)

**E**

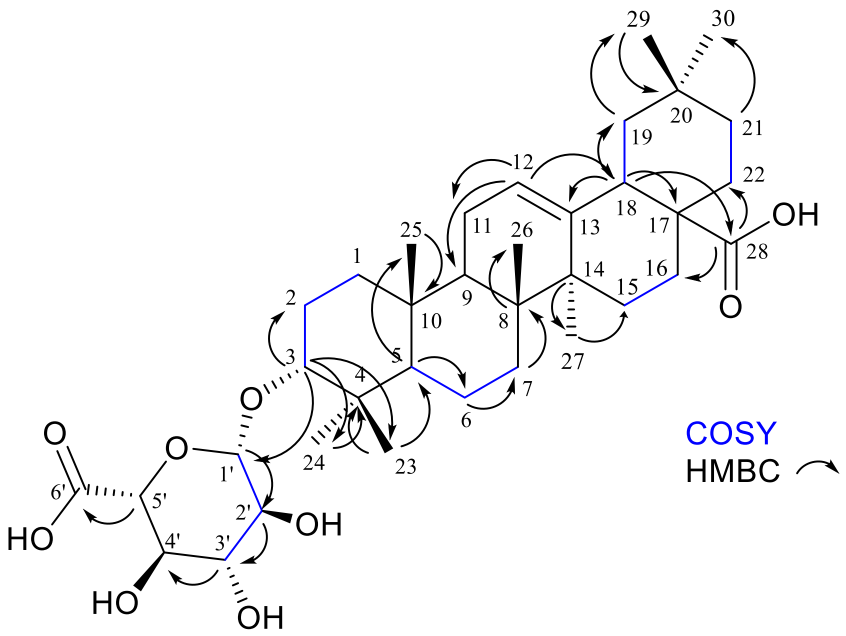


**Figure S11. Oleanolic acid-3-O-β-D-glucuronoside structural elucidation.**

The 1H NMR (**A**), 13C NMR (**B**), 2D COSY (**C**), and 2D HMBC (**D**) spectra of oleanolic acid-3-O-β-D-glucuronoside isolated from C. album line 3-4 seeds.

**E.** Key $^1H$–$^1H$ COSY (blue bonds) and HMBC (arrows) correlations used for the structure elucidation of oleanolic acid-3-O-β-D-glucuronoside.

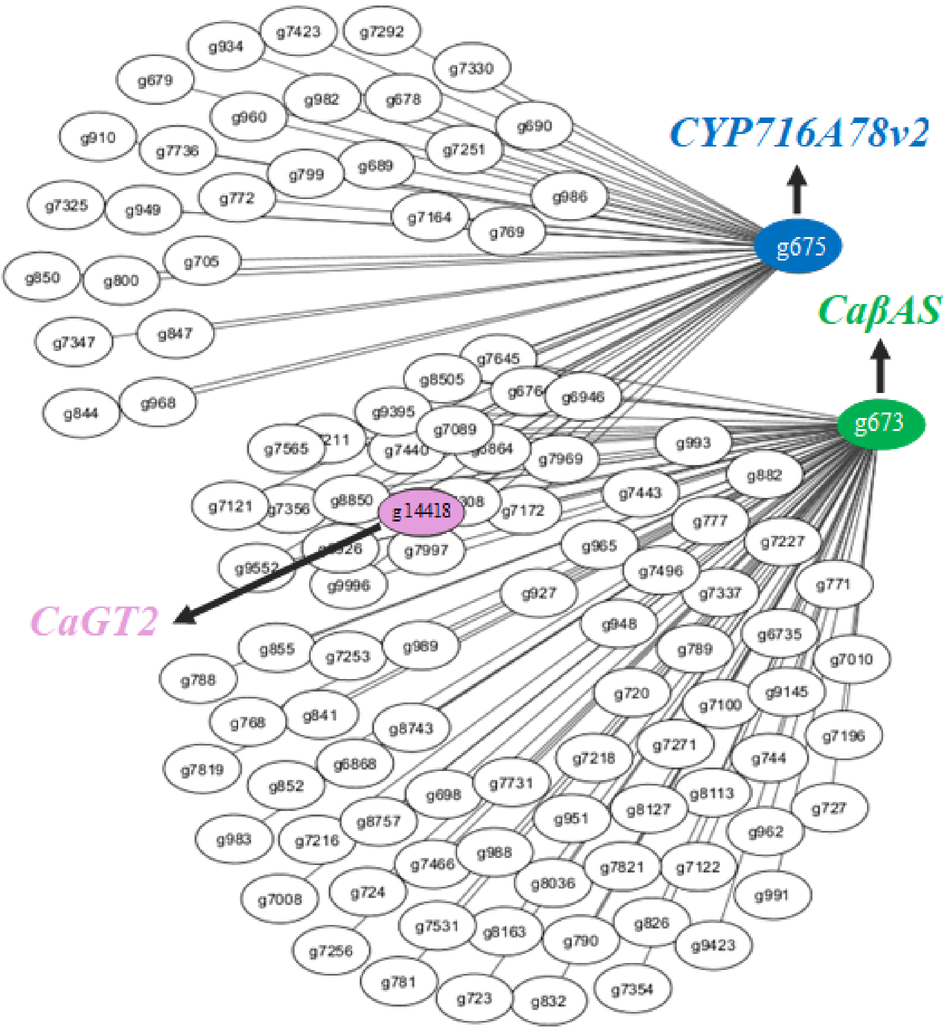


**Figure S12. Coexpression network of *C. album* genes associated with saponin biosynthesis.**

A gene coexpression network constructed using β-amyrin synthase (CaβAS; g673) and cytochrome P450 CYP716A78v2 (CaCYP716A78v2; g675) as bait genes is shown. Nodes represent genes, and edges indicate coexpression relationships based on transcriptomic data. Pearson pairwise correlation coefficient were used with a cutoff of r ≥ 0.8. In total 128 genes were coexpressed, including 80 genes correlated with g673 and 48 genes correlated with g675. The glucuronosyltransferase CaGlcAT2 (g14418) is highlighted within the network.

### Methods

#### Plant materials and growth conditions

A wild *Chenopodium album* core collection was established from seeds collected from individual plants across Denmark. Lines were grown in a climate-controlled greenhouse under long-day conditions (16 h light/8 h dark) at 21 °C and were self-pollinated. Propagation was carried out by single-seed descent up to the S5 generation (n = 4 per line) to reduce within-line heterogeneity. Two lines contrasting in seed saponin content were selected for detailed analyses: the high-saponin line 3-4 and the low-saponin line 6-1. For downstream experiments, plants were grown under the same greenhouse conditions, and leaves (third to fourth internode from the apex) and mature seeds were harvested at the S4 stage. Three biological replicates were collected per tissue and per line. This plant material was used for genome sequencing, RNA sequencing, metabolomic analyses, and quantification of seed protein and saponin content.

#### Total saponin and protein content

Seed saponin content of *C. album* lines was estimated using a foam test as previously described (Koziol, 1991). Briefly, 0.5 g of seeds were suspended in 5 mL of distilled water and agitated vigorously for 30 s, and foam height was recorded as a relative measure of total saponin content. For protein quantification, total nitrogen content was determined by CHNS elemental analysis. Exactly 25 mg of seed material was weighed and analyzed using a Vario MACRO Cube CHNS Elemental Analyzer (Elementar). A sulfanilamide standard was included as a reference, and nitrogen content was converted to protein content using a nitrogen-to-protein conversion factor of 6.25 (Food, 2003; Bonke et al., 2020).

#### Chromosome counting, ploidy analysis and genome size estimation

Chromosome counts were performed using young root tips collected from seedlings of ten *C. album* lines indicated in Table S6. Root tips were treated with 0.3% colchicine to arrest cells in metaphase, followed by fixation in ethanol:acetic acid (3:1, v/v). Fixed material was enzymatically digested using a cellulase and pectolyase mixture, and chromosome spreads were prepared using a steam-drop protocol (Kirov et al., 2014). Slides were stained with DAPI and imaged using a fluorescence microscope, and chromosome numbers were determined using DRAWID based on at least three well-spread metaphase images per line (Kirov et al., 2017).

Genome sizes were analyzed using a Quantum P flow cytometer equipped with a 488 nm laser (180 mW) and Cypad Software (Quantum Analysis) (Van Oost et al., 2021). Sample preparation was performed using the CyStain PI kit (Sysmex) according to the manufacturer's protocol. Fresh young leaf samples (±0.5 cm2) of the *C. album* plants and of the internal standard maize (Zea mays 'CE-777′: 5.43 pg/2C) were co-chopped using a razor blade in 0.4 mL of extraction buffer. A 50 µm CellTrics filter (Sysmex) was used to filter the chopped samples. Subsequently, the samples were stained in 1.2 mL staining buffer (CysStain PI kit) with propidium iodide as intercalating DNA stain. The samples were incubated in the dark at 4 °C for at least 30 minutes before analysis. Three replicate samples were prepared for each genotype on three different days from different plants of the same genotype with at least 3000 particles were analyzed per run. For each analysis two histograms were obtained (FL2 and FL3 detector). Genome sizes were calculated from the peak positions of the *C. album* nuclei and the internal reference *Zea may*s with known genome size: 2C = 5.43 pg. Mean values and standard deviations were calculated based on the three replicates and two histograms per replicate. Ploidy levels were analyzed as described in Akbarzadeh et al. (2025) but without the use of an internal standard. Leaf samples used and chopping was performed as described for the genome size analysis. The chopped samples were filtered through a 50 µm mesh nylon and 750 µl of the staining buffer was added (0.4 M Na2HPO4, 2 mg.L− 1 4′,6-diamidino-2-phenylindole (DAPI), pH 8.5). After staining the samples were analysed on a CyFlow Space flow cytometer (Sysmex) equipped with a 365 UV-LED. Histograms were plotted with fluorescence intensity on a linear scale (X-axis) and the number of DNA-stained nuclei (Y-axis) appearing as peaks on the histogram. The first sample analyzed with known diploid ploidy level was used as a reference and positioned at intensity 100. Samples with a peak positioned at intensity ±200 were considered to be tetraploid.

**Preparation of plant extracts and metabolite analysis**

Leaves and seeds from *C. album* lines 3-4 and 6-1 (S4) were ground in liquid nitrogen, and 100 mg of tissue was extracted with 300 µL of 85% (v/v) methanol. Samples were vortexed for 1 min, sonicated for 30 min, and centrifuged at 10,000 × g for 10 min, and the supernatant was filtered through a 0.22 µm membrane prior to analysis. LC–MS analysis was performed following the method described by Khakimov et al. (2015). Chromatographic separation was carried out on a Phenomenex Kinetex XB-C18 column at 40 °C with a flow rate of 0.3 mL/min, using water (A) and acetonitrile (B), both containing 0.1% formic acid, with a gradient from 5% to 100% B. The injection volume was 2 μL. Mass spectrometric analysis was conducted on a Bruker Daltonics Compact QqTOF mass spectrometer equipped with an electrospray ionization source operating in negative ion mode, acquiring MS/MS spectra over an m/z range of 50-1200. Data were calibrated using sodium formate clusters and processed with Bruker DataAnalysis software.

**Sample collection and DNA isolation**

For genome sequencing and resequencing, genomic DNA was isolated from leaf tissue of *C. album* lines 6-1 and 3-4, respectively. High-molecular-weight DNA for PacBio long-read sequencing was extracted from mature leaves of a single soil-grown plant of line 6-1 using a QIAGEN Genomic-tip (Cat no./ID. 10243), following the manufacturer's instructions. For Illumina short-read sequencing, genomic DNA was extracted from leaves of a single soil-grown plant of line 3-4 using the Qiagen DNeasy Plant Mini Kit (Cat no./ID. 69104). DNA quality and integrity were assessed by agarose gel electrophoresis and quantified using nanodrop and Qubit fluorometry.

**PacBio library preparation and sequencing**

A HiFi whole-genome shotgun library was prepared using the PacBio SMRTbell library preparation protocol and sequenced on a PacBio Sequel II platform, following the standard procedures for HiFi sequencing.

**Paired-end (PE) DNA library preparation and sequencing**

Genomic DNA from *C. album* line 3-4 was used for Illumina short-read resequencing. DNA quality and integrity were assessed using spectrophotometry and electrophoretic methods. Paired-end sequencing libraries were prepared following standard Illumina genomic DNA library preparation protocols and sequenced on an Illumina NovaSeq 6000 platform using a 2 × 150 bp configuration.

#### *De novo* genome assembly for tetraploid *C. album*

PacBio HiFi long-read data were used to generate a *de novo* genome assembly for *C. album* line 6-1, a low-saponin tetraploid line. The resulting assembly is referred to as Ca6-1v1. GenomeScope2 (v2.0) was applied to k-mer frequency profiles to obtain approximate estimates of genome characteristics (Ranallo-Benavidez et al., 2020). *De novo* assembly was performed using Hifiasm (v0.16) (Cheng et al., 2021). Assembly polishing and quality control were carried out in two steps: contigs were first polished using Racon (v1.4.3), and potential contaminant sequences were subsequently identified using Kraken2 (v2). Contigs classified as non-Viridiplantae were removed (Vaser et al., 2017; Wood et al., 2019), resulting in the exclusion of 18 contaminant contigs.

#### Quality assessment of the genome assembly

The quality of the Ca6-1v1 genome assembly was assessed using Benchmarking Universal Single-Copy Orthologs (BUSCO v5.7.0) and the QUality ASsessment Tool (QUAST) (Simão et al., 2015; Gurevich et al., 2013; Manni et al., 2021). BUSCO was used to evaluate assembly completeness based on conserved single-copy orthologs, using the Eukaryota, Viridiplantae, and Eudicots datasets. Assembly contiguity and general statistics, including total assembly length and scaffold number, were assessed using QUAST.

#### Annotation and gene prediction

The Ca6-1v1 genome assembly was soft-masked for transposable elements using EDTA (v1.9.6) (Ou et al., 2019). The soft-masked assembly was used as input for BRAKER (v3.0.8) (Hoff et al., 2019; Brůna et al., 2021), which integrates AUGUSTUS (v3.5.0) and GeneMark for gene prediction, incorporating RNA-seq evidence and a set of Viridiplantae protein sequences downloaded from UniProt to improve model training (Brůna et al., 2021). RNA-seq reads were trimmed and mapped to the reference genome prior to use in gene prediction. Functional annotation of predicted protein sequences was performed using the PANNZER2 webserver (v2.0) with a minimum query or subject coverage of 0.2 and a minimum alignment length of 40 (Törönen et al., 2018).

### Resequencing analysis and variant identification

Illumina short-read resequencing data were generated for *C. album* line 3-4 and used to identify sequence variation relative to the Ca6-1v1 reference genome. Reads were trimmed using Trimmomatic with a minimum read length of 35 bp and a minimum base quality of 20 (Bolger et al., 2014) and mapped to the Ca6-1v1 assembly using minimap2 (v2.17) (Li, 2018). Mapped reads were sorted and deduplicated prior to variant calling. Single-nucleotide variants and small insertions and deletions were identified using Platypus (v3), considering only uniquely mapping reads (MAPQ > 30). Variant calls were filtered using bcftools (v1.20) with a minimum variant quality of 30 and minimum read depth of 6 (Li, 2011). Functional effects of variants were annotated using Ensembl Variant Effect Predictor (VEP, release 112) with a custom *C. album* annotation file (McLaren et al., 2016). Structural variants, including deletions, insertions, inversions, and duplications, were identified using Manta based on paired-end and split-read evidence (Chen et al., 2016).

### RNA isolation and RNA-seq analysis

Total RNA was extracted from leaves and mature seeds of *C. album* lines 3-4 and 6-1 using the Spectrum™ Plant Total RNA Kit, followed by DNase treatment to remove genomic DNA. Three biological replicates were collected per tissue and per line, resulting in a total of 12 RNA-seq libraries. RNA quality was assessed prior to library preparation, and libraries were sequenced on an Illumina NovaSeq 6000 platform using paired-end sequencing. Raw reads were trimmed using Trimmomatic with a minimum read length of 35 bp and a minimum base quality threshold of 20 (Bolger et al., 2014). Transcript abundances were quantified using kallisto (v0.46) against the *C. album* transcriptome generated during genome annotation (Bray et al., 2016). Gene-level count matrices were constructed using the R package tximport and used for downstream analyses. Variance-stabilizing transformation was applied using DESeq2, followed by principal component analysis. Hierarchical clustering was performed using Euclidean distance as implemented in the pheatmap R package.

### Differential expression and gene enrichment analysis

Differential gene expression analyses were performed for the following comparisons: leaves of line 3-4 versus leaves of line 6-1, seeds of line 3-4 versus seeds of line 6-1, leaves versus seeds of line 3-4, and leaves versus seeds of line 6-1. Differential expression was assessed using DESeq2, and genes with an absolute log2 fold change greater than 1 and a false discovery rate (adjusted p value) below 0.05 were considered differentially expressed. Prior to analysis, low-count genes were filtered using the HTSfilter package. Gene Ontology enrichment analysis was conducted separately for up- and down-regulated genes in each comparison using the clusterProfiler package. For all enrichment analyses, categories with an adjusted p value below 0.05 and containing at least three differentially expressed genes were reported. In cases where no significantly enriched categories were identified, no Gene Ontology results are shown.

**Identification of saponin biosynthetic genes in *C. album***

Genes involved in triterpenoid saponin biosynthesis in *C. album* were identified using homology-based searches of the *C. album* line 6-1 proteome against a curated dataset of known saponin biosynthetic enzymes representing major gene families, including oxidosqualene cyclases (OSCs), CYP716 cytochrome P450s, and glucuronosyltransferases (GlcATs), from multiple plant species (Chung et al., 2020; Günther et al., 2022; Feng et al., 2024). BLASTp searches were performed using CLC Main Workbench. For each gene family, candidate homologs were ranked by sequence similarity, and the top hits with the lowest E-values and highest percentage identity were retained. Additional homologs of saponin biosynthetic genes were identified using *C. quinoa* genes as queries for RNA-seq-based allele mining, retaining the top five hits per gene (Tables S13, S14, and S16). Expression profiles of candidate genes were extracted from RNA-seq datasets generated for lines 3-4 and 6-1. Sequence variants were identified by intersecting gene coordinates with variant call files derived from Illumina resequencing data. The same gene sets were used for downstream phylogenetic analyses.

**Gene cluster analysis**

Biosynthetic gene clusters in *C. album* were predicted using plantiSMASH, with the Ca6-1v1 genome assembly and corresponding gene annotation provided as input (Kautsar et al., 2017).

**Gene cloning and tobacco infiltration**

Coding sequences for oxidosqualene cyclase/β-amyrin synthase (OSC/CaβAS; g673), three cytochrome P450 enzymes (CYP716A78v1, g4225; CYP716A78v2, g675; CYP716A78v3, g33569), and two glucuronosyltransferases (GlcATs; CaGlcAT1, g2573; CaGlcAT2, g14418) were amplified from cDNA derived from *C. album* seeds using gene-specific primers (listed in Table S19). PCR products were gel purified and initially cloned into the pJET1.2 blunt-end cloning vector. Inserts were subsequently subcloned into the pEAQ-HT-DST1 expression vector using NEBuilder HiFi DNA Assembly. Final constructs were transformed into *Agrobacterium tumefaciens* strain AGL1.

For transient expression, *N. benthamiana* plants were grown under controlled conditions (24 °C, 16 h light/8 h dark). Agrobacterium cultures carrying individual or combined constructs were adjusted to an $OD_{600}$ of 0.5 in infiltration buffer (¼ MS, 1% sucrose, 100 μM acetosyringone, 0.005% Silwet L-77) and infiltrated into leaves by syringe. CaβAS was expressed alone or co-expressed with individual CYP716 enzymes, followed by co-expression with GlcATs. Wild-type plants, buffer-only infiltrations, and RUBY-expressing plants used as visual infiltration control were included. Each treatment was performed with three biological replicates. Leaf tissue was harvested 5 days post infiltration, flash-frozen, and stored at −80 °C prior to GC–MS and LC–MS analysis.

**GC–MS analyses for β-amyrin and oleanolic acid quantification**

Leaf tissue from wild-type and infiltrated *N. benthamiana* plants was used for GC–MS analysis (Liu et al., 2019). Samples (150 mg) were ground in liquid nitrogen and extracted with ethyl acetate, followed by sonication for 30 min. After centrifugation, supernatants were filtered and derivatized with trimethylsilyl cyanide for 50 min at 70 °C. GC–MS analysis was performed on a Shimadzu GC-2010 system coupled to an MS-QP2010 Plus detector. Compounds were separated on an HP-5MS column (30 m × 250 μm × 0.25 μm) using helium as the carrier gas at a constant flow rate of 11.9 mL $min^{-1}$. The oven temperature program was as follows: 60 °C for 1 min, ramped at 30 °C $min^{-1}$ to 280 °C and held for 2 min, followed by a ramp of 1 °C $min^{-1}$ to 310 °C and a final hold of 4 min. Mass spectra were acquired in the range of 50–600 m/z. Transfer line, ion source, and quadrupole temperatures were set to 280 °C, 230 °C, and 150 °C, respectively. GC–MS data were processed using GCMSsolution (v4.2, Shimadzu) software. β-Amyrin and oleanolic acid were identified based on retention times and fragmentation patterns compared with authentic standards, and relative abundances were estimated by extracting characteristic fragment ions (m/z 129, 203, 218, 320, and 471).

**Saponin purification and NMR analysis**

Oleanolic acid monoglucuronoside was purified from seeds of *C. album* line 3-4. Approximately 500 g of seeds were ground and extracted with 80% (v/v) ethanol. The crude *C. album* seed extract was further purified via a PuriFlash 5.250 coupled with an ELSD detector using a Uptisphere® StrategyTM PF-15C18HQ-F0120 Interchim® column (100 Å, 15 µm C18, 224 x 36 mm, Interchim, Montluçon, France). The purified fraction was analyzed by Agilent 1260 Infinity II LC/MSD system to confirm its identity based on m/z values compared with those detected in crude seed extracts. For HPLC analysis and preparative purification, high purity Milli-Q grade water and acetonitrile (LC-MS grade, VWR) and both solvents were supplemented with 0.1% (v/v) formic acid (98%-100%, LiChropur, Sigma) were employed as mobile phases A and B, respectively. Gradient conditions were as follows: 0.0–1.0 min 5% B; 1.0–20.0 min 5%–60% B, 20.0–25.0 min 20%–95% B, 25.0–30.0 min 95% B, 30.0–35.0 min 5% B and 35–38.0 min 5% B.

Initial compound detection system optimization was performed with the Flash purified *Chenopodium album* extract F2 using a ZORBAX SB-C18 analytical column (150 mm × 4.6 mm i.d., 5 µm particle size, 100 Å pore size;). The flow rate was set at 0.5 mL/min. Mass spectrometric analysis was carried out in negative ionization mode. Both full-scan and Selective Ion Monitoring (SIM) were used, targeting ions at m/z 455, 631, and 677, which correspond to diagnostic fragments of the compound of interest. Instrument-specific ion optics, source parameters, and acquisition settings were kept consistent across runs for comparative integrity. The MSD spray chamber was equipped with a Agilent Electiospray Ionization source. The capillary voltage was 3000 V, Drying Gas Flow was 12 L/min (nitrogen), Nebulizer Pressure was 35 psig, and the drying gas temperature was 350 °C. The fragmentor energy was set to 70 eV, 62 msec Dwell time for each SIM fragment and MS scan was performed from mass 450 – 1300 with fragmentor set to 70 eV and 1733 cycle speed (units/ sec). Purification of the target compound was carried out using the same system in preparative mode, with a Kinetex XB-C18 semi-preparative column (150 mm × 10 mm i.d., 5 µm, 100 Å; Phenomenex). The mobile phase and gradient were identical to the analytical method, scaled for a flow rate of 4.0 mL/min. Detection was performed using the Variable Wavelength Detector (VWD) set to 205 nm. Fractions were collected based on a time-based window corresponding to the UV-detected peak: 22.0 to 24.9 minutes. Collected fractions were pooled and concentrated under reduced pressure using a rotary evaporator and subsequently lyophilized. Saponins were analyzed on a Dionex UltiMate 3000 UHPLC system (Thermo Fisher Scientific, Germany) coupled to a Bruker Compact qToF-MS with ESI source (Bruker Daltonics, Germany). Separation was achieved on a Kinetex 1.7 µm XB-C18 column using high purity Milli-Q grade water and acetonitrile (both solvents were supplemented with 0.05% (v/v) formic acid) were employed as mobile phases A and B, respectively. Flow rate was 300 µL/min, column temperature was 30°C (Trinh et al., 2024).

Approximately 6 mg of the purified compound was dissolved in DMSO-$d_6$ (99.5%, isotopic) and analyzed at 25 °C using a Bruker Avance Neo 800 MHz spectrometer equipped with a 5 mm CPTXO Cryoprobe. Data acquisition and processing were conducted using TopSpin v4.07 (Bruker). Spectral figures were prepared using MestReNova sorftware (Mnova, Mestrelab Research).

### Data S1. Sequences of *C. album* genes cloned in this study.

> CaβAS_g673
ATGTGGAGGTTAAAAGTTGGGGAAGGTGCAAATGACCCTTATTTATACAGCACTAATAACTTTGTTGGACGACAAACTTGGGA
GTTTGATCCTAACTATGGCACCCCTGAGGAGAGGGAGGAGGTGGAGGAAGCACGTCGCAACTTCTACAACAATCGATTTAAAG
TTAAGCCTTGTGGTGATCTCATATGGCGTCTTCAGTTCCTAAGGGAGAAAAACTTCAAGCAAACAATACCTCAAGTGAAGGTG
GAAGAAGGGGAGGAGATCACATACGAAACCGCGACGACAACATTAAAGAGAGCCGTGAATGTATTCACAGCCCTGCAGTCTG
ACCAAGGCCATTGGCCTGCTGAAATTGCTGGCCCTCAATTTTTCCTTCCTCCCTTGGTTTTCTGCTTATACATTACAGGGGATCT
GAATTCTGTTTTCGGGCCAGAGCATCGTAGAGAGATTCTTCGTAGTATCTATTATCATCAGAATGAAGATGGAGGTTGGGGATTG
CATATTGAAGGACACAGTACCATGTTTTGTACTGCACTCAACTATATTTGTTTGCGAATGCTTGGAATCGGACCGGATGAGGGTG
ATGACAATGCTTGTCCTAGGGCTCGAAAATGGATCCTTGACCATGGTAGTGTTACCCATATGCCTTCTTGGGGAAAAACTTGGC
TTTCTATACTCGGCTTGTTTGATTGGTCTGGAAGTAATCCAATGCCACCTGAGTTCTGGCTCCTTCCGTCTTTTCTCCCTATGTAT
CCTGCAAAAATGTGGTGCTATTGTCGAATGGTATACATGCCAATGTCATATTTGTATGGGAAGAGATTTGTAGGTCCGATCACAC
CTCTCATTAAACAACTTAGAGAAGAACTCTACAACGAACCCTTTGAGCATATTAGTTGGAAGCAAATGCGTCATTTGTGTGCAC
CGGAGGATCTTTACTACCCTCATCCATTGATTCAAGATTTAATGTGGGACGCTCTTTACATCTTTACGGAGCCTCTCCTTACTCGT
TGGCCTTTCAACAAGCTGATTCGAAAGAAGGCACTCGAGGTTACAATGGAACACATTCATTACGAAGATGAGAACAGTCGTTA
CATAACCATTGGAAGTGTTGAGAAGGTTTATGCATGTTAGCTTGTTGGGTTGAAGACCCTAAAGGAGATCATTACAAGAAACA
TCTTGCTAGAGTCCAAGATTACATTTGGATTGCTGAAGATGGATTAAAAATGCAGAGTTTTGGAAGTCAAGAATGGGATTGTGG
GTTTTCCGTCCAAGCATTGCTAGCTTCAAATCTTAGCCTCGACGAAATTGGACCTGCTCTTAAGAAAGGCCACTTCTTTATTAAA
GAGTCACAGGTTAAGGACAATCCATCAGGCGACTTCAAAGCTATGCACCGCCATATCTCAAAGGGATCATGGACTTTTTCCGAC
CAAGATCATGGTTGGCAAGTCTCTGATTGTACCGCGGAAGGCCTTAAGTGTTGTCTAATCTTATCGACAATGCCCCCGGAAATT
GTTGGAGAAAAGATGGACCCTGAACGACTTTATGATTCAGTCAATGTCTTACTTTCTCTACAGAGTAAAAATGGCGGGCTAGCT
GCTTGGGAACCTGCAGGGGCTCAAGAATGGTTGGAGGTTCTTAACCCAACAGAATTTTTTGAAGGCATTGTCATTGAATATGA
GTACGTAGAATGTACAGCTTCAGCAATTCAAGCTCTAGTTATGTTCAAGAAGTTATACCCAGGCCACAGGAAGAAAGAGATCG
ATAATTTCGTTGTAAATGCAGTCCGGTACCTTGAGAACACCCAGTTTCCTAGTGGAGGATGGTATGGAAATTGGGGTATTTGCTT
CATATATGGGACATGGTTTGCTCTTGGAGGGCTAGCAGCTGGTGGGAAAACGTACTACAATTGTGCTGCTGTTAGGAAGGGTGT
CGAATTTCTACTTACAACGCAAAAGGAGGATGGTGGATGGGGTGAGAGTTATATTTCGTGTCCCAAAAAGGAATTTGTGCCAAT
AGAAGGAAAGTCCAATTTGGTCCAAACCGGATGGGCTTTAATGGGTCTACTTCATGCTGGACAGGCGGAGAGGGATCCAACTC
CTCTACATCGTGCAGCAAAGCTTTTGATCAACTCACAACTCGAAAATGGTGATTTTCCCCAACAGGAAATAACAGGGGTGTTC
ATGAAGAATTGCATGTTACATTATCCAATGTATAGGAGTATTTACCCAATGTGGGCACTTGCGGAGTACAGAAAGCGTGTTTCAT
TACCTTCTATCAACTCTGCCTAA

>CaCYP716A78v1_g4225
ATGGAGCTCTTCTTCTTGGCTAGCATAGCCCTTATTTCCTCCTTATCTCTACCTTTTCTCTATCTATTCTATAGGCATTATTCGACCT
GGGGGTACAAGCTACCCCCAGGATCGATGGGATGGCCCGTGGTGGGCGAATCCCTAGAGTTTTTCTCTACCGGTTGGAAGGGA
TACCCAGAGAAGTTCATCTTTGATAGACTTAAAAAGTACAAACCTAGCCAAGTGTTCAAGACTTCCATCTTTGGTGAAAAGGTT
GCAATTTTATGTGGCGCGACAGGTAACAAATTCTTGTACTCGAACGAGAACAAGTTAGTACAAGCTTGGTGGCCTAAATCAGTT
GACAAGATCTTTCCTGCTGCCACCCAACATTCCTCCATAGAAGAAGCTAGGACTATGCGGAAGCTTATCCCTCTTTTCCTTAAG
CCCGAATCTTTACAAAGGTACATACCCATCATGGACACCATAGCCACCAGGCACATGGAGTCCGGGTGGGAGGGCAAGGACAA
GGTAGAAGTGTTCCCATTAGCTAAGCGATACACCTTTTGGGTGGCTTGTAGGCTCTTCTTGAGCATTGAGGACCCGGTCCATGT
AGCCAAGTTCGCGGACCCCTTCAATGAGATAGCCGCAGGGATCATATCCATCCCAATAGATCTCCCCGGGACACCGTTCCACAA
AGGGACCAAATCTTCTGAAATCGTAAGGAAAGAGTTGAGAGCAATTATCAAGCAAAGGAAATTAGACTTAGCAGATGGCAAA
GCTTCACCTACACAAGATATTCTATCTCATATGTTGTTAACTTCTACTGATGATGGGAAGTTTATGAATGAAATGGATATTGCTAAT
AAAATTTTGGGACTTCTTATTGGTGGACATGATACTGCTAGTGCTTCTTGTACCTTTATTGTCAAGTTTCTTGCTGAACTTCCTCA
CATCTATGAAGGTGTTTACAAAGAGCAAATGGAGATAGCAAATTCAAAAAAACCAGGGGAGCTTCTAAATTGGGAGGACATAC
AAAAGATGAAATACTCATGGAATGTGGCATGTGAAGTGATGCGTTTGGCTCCTCCACTCCAAGGTGGTTTTAGAGAAGCCATTT
CCGACTTTATGTACAACGGATTCCAAATTCCCAAGGGCTGGAAGCTATATTGGAGTGCAAATACAACACATTTGAACCCAGAAT
GTTTCCCGGAACCAACGAAATTCGACCCATCGAGGTTCGATGGGTCCGGGCCAGCACCATACACATTCGTACCCTTTGGAGGG
GGACCAAGAATGTGCCCAGGAAAAGAGTATGCAAGGCTAGAGATATTGGTGTTCATGTACCATATTGTCAAGAGGTTTAAATGG
GAAAAAGTGCTTCCTACTGAGAAAGTTATTGTTAATCCCATGCCTATTCCGGAGCACGGCCTTCCCGTCCGCCTTTTCCCACATC
CTCAAACCACGGTTGCTTAA

> CaCYP716A78v2_g675
ATGGAGCTCTTCTTCTTGTATAGCATAGCCCTTATTTCCTCCTTATCTCTACCTTTTCTCTACCTATTCTATAGGCATTATTCGACCT
GGGGGTACAAGCTACCCCCAGGGTCGATGGGATGGCCCGTGGTGGGTGAGTCCCTAGAGTTTTTCTCTACCGGTTGGAAGGGA
TACCCGGAGAAGTTCATCTTTGATAGACTTAAGAAGTACAAACCTAGCCAAGTGTTCAAGACTTCCATCTTTGGTGAAAAGGTT
GCGATTTTATGTGGAGCGACAGGTAACAAGTTCTTGTACTCGAACGAGAACAAGTTAGTACAAGCTTGGTGGCCTAAATCAGT
TGATAAGATCTTTCCTGCTGCCACCCAACATTCCTCCATAGAAGAAGCTAGGACTATGCGGAAGCTTATCCCTCTCTTCCTTAAG
CCCGAATCTTTACAAAGGTACATACCCATCATGGACACCATAGCCACCAGGCACATGGAGTCCGGGTGGGAGGGCAAGGACAA
GGTAGAAGTGTTCCCCTTAGCAAAGCGATACACCTTTTGGGTTGCTTGTAGGCTCTTCTTGAGCATCGAGGACCCGGTCCATGT
AGCCAAGTTCGCCGACCCCTTCAATGAGATAGCCGCAGGGATCATATCCATCCCAATAGATCTCCCCGGGACACCGTTCCACAA
AGGGATCAAATCTTCCGAAATCGTAAGGAAAGAGTTGAGAGCAATTATCAAGCAAAGGAAATTAGACTTAGCAGATGGCAAA
GCTTCACCAACACAAGATATTCTATCTCATATGTTGTTAACTTCTACCGATGATGGGAAGTTTATGAATGAAATGGATATTGCTAA
TAAAATTTTGGGACTTCTTATTGGTGGACATGATACTGCTAGTGCTTCTTGTACCTTTATTGTCAAGTTTCTTGCTGAACTTCCTC
ACATCTATGAAGGTGTTTACAAAGAGCAAATGGAGATAGCTAATTCAAAAAAACCAGGGGAGCTTCTAAATTGGGAGGACATT
CAAAAGATGAAATACTCATGGAATGTGGCATGTGAAGTGATGCGTTTGGCTCCTCCACTCCAAGGTGGTTTTAGAGAAGCCATT
TCCGACTTTATGTACAACGGGTTCCAAATTCCCAAGGGCTGGAAGCTATATTGGAGTGCAAATACAACACATTTGAACCCAGAA
TGTTTCCCGGAACCAACGAAATTCGACCCATCGAGGTTCGATGGGTCCGGGCCAGCACCATACACATTTGTACCCTTTGGAGG
GGGACCAAGAATGTGCCCAGGAAAGGAGTATGTAAGGCTAGAGATATTGGTGTTCATGTACCATATTGTCAAGAGGTTTAAATG
GGAAAAAGTGCTTCCTACTGAGAAAGTTATTGTTAATCCCATGCCTATTCCGGAGCACGGCCTTCCCGTCCGCCTTTTCCCACAT
CCTCATACCACGGTTGCTTAA

> CaCYP716A78v3_g33569
ATGGAGCTCATCTTCTTGGCTAGCATAGCCCTTATTTCCTCCTTATCTCTACCTTTTCTCTATCTATTCTATAGGCATTATTCGACCT
GGGGGTACAAGCTACCCCCAGGATCGATGGGATGGCCCGTGGTGGGCGAATCCCTAGAGTTTTTCTCTACCGGTTGGAAGGGA

```
TACCCAGAGAAGTTCATCTTTGATAGACTTAAAAAGTACAAACCTAGCCAAGTGTTCAAGACTTCCATCTTTGGTGAAAAGGTT
GCAATTTTATGTGGCGCGACAGGTAACAAATTCTTGTACTCGAACGAGAACAAGTTAGTACAAGCTTGGTGGCCTAAATCAGTT
GACAAGATCTTTCCTGCTGCCACCCAACATTCCTCCATAGAAGAAGCTAGGACTATGCGGAAGCTTATCCCTCTTTTCCTTAAG
CCCGAATCTTTACAAAGGTACATACCCATCATGGACACCATAGCCACCAGGCACATGGAGTCCGGGTGGGAGGGCAAGGACAA
GGTAGAAGTGTTCCCATTAGCTAAGCGATACACCTTTTGGGTGGCTTGTAGGCTCTTCTTGAGCATTGAGGACCCGGTCCATGT
AGCCAAGTTCGCGGACCCCTTCAATGAGATAGCCGCAGGGATCATATCCATCCCAATAGATCTCCCCGGGACACCGTTCCACAA
AGGGATCAAATCTTCTGAAATCGTAAGGAAAGAGTTGAGAGCAATTATCAAGCAAAGGAAATTAGACTTAGCAGATGGCAAAG
CTTCACCTACACAAGATATTCTATCTCATATGTTGTTAACTTCTACTGATGATGGGAAGTTTATGAATGAAATGGATATTGCTAATA
AAATTTTGGGACTTCTTATTGGTGGACATGATACTGCTAGTGCTTCTTGTACCTTTATTGTCAAGTTTCTTGCTGAACTTCCTCAC
ATCTATGAAGGTGTTTACAAAGAGCAAATGGAGATAGCAAATTCAAAAAAACCAGGGGAGCTTCTAAATTGGGAGGACATACA
AAAGATGAAATACTCATGGAATGTGGCATGTGAAGTGATGCGTTTGGCTCCTCCACTCCAAGGTGGTTTTAGAGAAGCCATTTC
CGACTTTATGTACAACGGATTCCAAATTCCCAAGGGCTGGAAGCTATATTGGAGTGCAAATACAACACATTTGAACCCAGAATG
TTTCCCGGAACCAACGAAATTCGACCCATCGAGGTTCGATGGGTCCGGGCCAGCACCATACACATTCGTACCCTTTGGAGGGG
GACCAAGAATGTGCCCAGGAAAAGAGTATGCAAGGCTAGAGATATTGGTGTTCATGTACCATATTGTCAAGAGGTTTAAATGGG
AAAAAGTGCTTCCTACTGAGAAAGTTATTGTTAATCCCATGCCTATTCCGGAGCACGGCCTTCCCGTCCGCCTTTTCCCACATCC
TCAAACCGCGGTTGCCTAA

>CaGlcAT1_g2573
ATGGGGGCAACACACATTTGCAAAGTCCAAACCAAAAGAGCCATTATCAACCGTATTCACATCCTCATACACTCCTTTGCCATT
CTTGCTCTCTTCTACTACCGTTTTTCGTCTTTCTCCAACCCTCATATCTCCCTTCTTCCATGGATATTATTGACCATCGCCGACCTC
GTTTTCACCTTCATTTGGGCCATGACCCAGGCCTTCCGTTGGCGCCCTGTCTTGCACGACGTGTCTGGCTATGAGTCCATCAATC
CACGTGATCTTCCAAAGATCGACATTTTTATATGCACCGCCGATCCCACAAAGGAGCCTGTGTTGGAAGTGATGAACTCGGTGA
TATCATCCATGGCGCTCGATTACCCGCCTGAAAAGATGGCGATATATTTGTCGGATGATGGAGGTTCTCCTTTGACTAGAGAGGC
TATTAAGAAGGCCGTTGAATTTGCTAAGGTTTGGATTCCTTTTTGTAACATGTATGCTATTAAGACTAGGTGCCCTGATGCTTTCT
TTTCCGCTTTGGGTGATGATGAAAGACTTCATCGTGATCAAGACTTTGACGCTCATGAATCACTGCTTAAGTCGAAATATGAAG
CTTTTAAGAAACATGTGGAGAAAGAAAGCGGTGATAATAATAAATGCACCGTTGTGCATGATCGTGCCCCTTGCATTGAGATTA
TACATGACAGTAAACAAGATGGAGAAGGAGAAGTGAAAATGCCCCTTGTAGTTTATGTAGCCAGGGAAAAGAGACCAGGTCA
TCCTCATCGTTTCAAAGCTGGAGCCCTTAACGCTCTTCTCCGAGTATCAGGACTATTGAGCAATGCGCCTTACTTATTGGTGTTG
GATTGTGATATGTACTGTCATGATCCAACCTCTGCTCGTCAATCTATGTGCTTCCATCTTGACCCTAACATGGCTCCTTCTCTTGC
CTTTGTTCAATACCCTCAAATTTCTACAACACGAGCAAAAATGATATCTATGATGGCCAAGCCAGATCAGCTCATACGACGAA
ATGGCAAGGCATGGATGGACTCAGAGGACCGGTCTTGAATGGAACTGGGTATTATCTGAAGAAGAAGGCGATATATGGAAGGC
CTCATAATGAAGATGAATACCTCATCAATGAACCAGAGAAGGCCTTTGGTTCTTCTACAAAATTCATCGCTTCACTTAAAGAAA
ACTCGAACCAGGATCTTGTCTTGAAGGAATTCACAAACGATTTGTTACAAGAGGCTAGAAATTTGGCTACTTGCACTTATGAAG
CAAACTCGCTATGGGGTGTTGAGGTAGGATTTTCGTATGATTGCCTGTTGGAGAGTTCATACACTGGATATCTCTTACATTGTAA
AGGATGGAGATCTGTGTATCTTTATCCCAAAAGACCGTGCTTCTTGGGATGCACGACAATTGACATGAAGGATGCTATTGTTCA
ATTAATAAAATGGACCTCCGGATTACTTGGAGTTGCCATGTCAAAGTTTAGCCCTCTTACCTATGCCATGTCCAGGATGTCTATTT
TGCAAAGCCTGTGTTATGCGTACATCACGTGTTCAGGTCTTCTTGCAGTTCCGCTCTTTATATATGGTGTTGTTCTTCCATTCTGC
CTACTTAAGGGCGTTCCTGTTTTTCCTAAGGTATCGGATCCATGGATGTTGGGTTTCGTGTTTGTATTTGTATCCTCCCATGTTCA
ACATCTATTCGAAGTGCTGGCAAGTGATCATTCAGTGCAACAGTGGTGGAATGAGGTGAGAATCTGGATCATGAAAGCGATAA
CAGCCTGCTTGTTTGGATCAACGGAAGCAGTAATGAAGAAGATTGGGATACAGAAAACAACATTCAGATTAACAAATAAGGTT
GTCGAGAAAGAGAAGTTGGATAAATACGAGAAGGGAAAGTTCGATTTCTCAGGGGCAGCAATGCTAATGGTTCCTCTCATCAT
TTTGACAGTACTAAATTTGGTGTCGTTCGTTGGGGGAATTATAAGGGTGATCAACCACAACAACTATGATGATATGTTTGGGCAA
CTTTTCCTGTCCTTTTATCTCCTGCTTCTTAGCTACCCTACTTTCGAAGGGATTGTTACAAAAATTACAGACAAACTTAGAAAGA
AAGAATAA

>CaGlcAT2_g14418
ATGGCGGCAACACACATTTGCAAAGTCCAAACCAAAAGAGCCATTATCAACCGTATTCATATCCTCATTCACTCTCTAGCCATTC
TTGCTCTCTTCTACTACCGTTTCTCGTCTTTCTCCAACCCTCATATCTCCCTTTTTCCATGGGTATTATTGACTATCGCCGACCTCG
TTTTCACCTTCATTTGGGCCATGACTCAGGCCTTCCGTTGGCGCCCCGTCTTGCACGATGTGTCTGGCTATGAGTCCATCAATCC
ACGCGATCTTCCAAAGATCGATATTTTTATATGCACCGCTGATCCCACCAAGGAGCCTGTGTTGGAAGTGATGAACTCGGTGAT
ATCATCCATGGCGCTCGATTATCCGCCTGAAAAGATGGCGGTGTATTTGTCGGATGATGGTGGTTCTCCTTTGACTAGAGAGGCT
ATTAAGAAGGCTGTTGAATTTGCTAAGGTTTGGATTCCTTTTTGTAATATGTATGCTATTAAGACTAGGTGTCCTGATGCTTTCTT
CTCCGCTTTGGGTGATGATGAAAGACTTCATCGTGATCAAGACTTTAACGCTCAGGAATCACTCCTCAAGTCGAAATACGAAGC
TTTTAAGAAATATGTGGAGAAAGAAAGCGGTGATATTAATAAATGCACCGTTGTGCATGATCGTGACCCTTGCATTGAGATTATA
CATGACAGTAAACAGGATGGAGAAGGAGAAGTGAAAATGCCCCTTGTAGTTTATGTAGCCAGGGAAAAGAGACCAGGTCATC
CTCATCGTTTCAAAGCTGGAGCCCTTAACGCTCTTCTCCGAGTATCAGGTCTATTGAGCAATGCGCCTTACTTATTGGTGTTGGA
TTGTGATATGTACTGTCATGATCCAACCTCTGCTCGTCAATCTATGTGCTTCCATCTTGACCCGAACATGTCTCCCTCTCTTGCCT
TTGTTCAATACCCTCAAATTTCTACAACACTAGTAAAAATGATATCTATGATGGTCAAGCCAGATCAGCTCATACGACGAAATG
GCAAGGCATGGATGGACTCAGAGGACCGGTCTTGAATGGAACTGGGTATTATCTGAAGAAGAAGGCGATATATGGAAGGCCCC
ATAATGAAGATGAATACCTCATCAATGAACCAGAGAAGGCCTTTGGTTCTTCCACAAAATTCATCACTTCACTCAAAGAAAACT
CCAACCAGGATCTTGTCTTGAAGGAATTCACAAACGATTTGTTACAAGAGGCTAGAAATTTGGCTACTTGCACTTATGAAGCAA
ACTCGCTATGGGGTGTTGAGGCAAGATTTTCGTATGATTGCCTGTTGGAGAGTTCATACACTGGATATCTCTTACATTGTAAAGG
ATGGAGATCTGTGTATCTTTATCCCAAAAGACCGTGCTTCTTGGGATGCACGACAATTGACATGAAGGATGCAATTGTTCAACT
AATAAAATGGACTTCCGGATTACTTGGAGTTGCCATGTCAAAGTTTAGCCCTCTTACTTATGCCATGTCCAGAATGTCTATTTTGC
AAAGCATGTGTTACGCGTACATCACGTGTTCAGGTCTTCTTGCAGTTCCACTCTTTATATATGGTGTTGTTCTTCCATTCTGCCTA
CTCAAGGGCATTCCTGTTTTTCCTAAGGTATCGGATCCATGGATGTTGGGTTTCGTGTTTGTATTTGTATCCTCCCATGTTCAACA
TCTATTCGAAGTGCTAGCAAGTGATCATTCAGTGCAGCAGTGGTGGAATGAGGTGAGAATCTGGATCATGAAAGCGATAACAG
CCTGCTTGTTTGGATCAACTGAAGCAATAATGAAGAAGATTGGGATACAGAAAACAACATTCAGATTAACAAATAAGGTTGTG
GAGAAAGAGAAGTTGGATAAATACGAGAAGGGAAAGTTCGATTTCTCAGGAGCAGCAATGCTAATGGTTCCTCTCATCATTTT
GACTATACTAAATTTGGTGTCGTTCGTTGGGGGACTTGTAAGGGTGATCAACCACAACAACTACGATGATATGTTCGGGCAACT
TTTCTTGTCATTTTATCTCCTACTTCTTAGCTACCCTACTTTCGAAGGGATTGTTACAAAAGTTACAGACAAACTTAGAAAGAAG
GAATAA
```
